# Epistasis of two classical color genes, *B* and *L-2*, synergistically controls carotenoid accumulation in squash

**DOI:** 10.64898/2026.05.19.726227

**Authors:** Lilin Xu, Xuesong Zhou, Emalee Wrightstone, Kay McNeary, Gregory Inzinna, Chris Hernandez, Zhangjun Fei, Harry S. Paris, Amit Gur, Arthur A. Schaffer, James Myers, Lailiang Cheng, Li Li, Michael Mazourek

**Affiliations:** Horticulture Section, School of Integrative Plant Science, Cornell University, Ithaca, New York 14853, USA; Plant Breeding and Genetics Section, School of Integrative Plant Science, Cornell University, Ithaca, New York 14853, USA; Robert W. Holley Center for Agriculture and Health, USDA-ARS, Cornell University, Ithaca, New York 14853, USA; Department of Agriculture, Nutrition, and Food Systems, University of New Hampshire, Durham, New Hampshire 03824, USA; Boyce Thompson Institute, Cornell University, Ithaca, 14853, USA; Department of Vegetable Sciences, Agricultural Research Organization, Newe Ya‘ar Research Center, Ramat Yishay, Israel; Institute of Plant Sciences, Agricultural Research Organization, The Volcani Center, Israel; Department of Horticulture, Oregon State University, Corvallis, OR 97331, USA

**Keywords:** *Cucurbita pepo*, carotenoids, epistasis, *B*, *L-2*, *PSY*

## Abstract

Carotenoid accumulation underlies fruit color and nutritional quality in squash (*Cucurbita pepo*). One pair of dominant genes, *B* and *L-2*, have been long known to interact epistatically, substantially boosting carotenoid accumulation and producing intensely orange-fleshed fruit. However, their molecular identities and regulatory mechanism are unknown. Here, we show that *B* encodes a truncated H subunit of magnesium chelatase (CpCHLH^B^) and *L-2* encodes a homolog of Arabidopsis Pseudo-Response Regulator 2 (CpAPRR2-A). Significantly, expression of *phytoene synthase* (*CpPSY-A*), which encodes the major rate-limiting enzyme in carotenoid biosynthesis, was dramatically upregulated in fruit of *B/B L-2/L-2* plants compared with *b/b L-2/L-2* or *B/B l-2/l-2*, showing that the *B* and *L-2* interaction affects *CpPSY-A* transcription. A similar upregulation was also observed in Arabidopsis *gun5 L-2* transgenic plants, where *gun5* is a genetic mimic of the *C. pepo B* gene. The wild-type CpCHLH^b^ physically interacted with CpAPRR2-A, attenuating the CpAPRR2-A-mediated activation of *CpPSY-A*. In contrast, the truncated CpCHLH^B^ lost its ability to interact with CpAPRR2-A, enabling CpAPRR2-A to activate *CpPSY-A* and produce intensely orange fruit. These findings uncover the mechanism underlying the epistatic interaction through which *B* and *L-2* act synergistically to boost carotenoid production, offering novel mechanistic insights and key targets for improving crop quality.

**One-sentence summary:** Synergistic epistasis between *B* and *L-2* arises from loss of interaction between their encoded proteins, resulting in dramatically upregulating the key rate-limiting enzyme in carotenoid biosynthesis pathway to produce intensely orange-fleshed fruit in squash.

## Introduction

The vibrant yellow, orange, and red colors seen in many food crops and ornamentals result from accumulation of carotenoids (Yuan et al. 2015; Hermanns et al. 2020). Beyond their visual impact, carotenoids play vital roles in plants as photosynthetic pigments and photoprotective agents. They also serve as precursors for the synthesis of the phytohormones abscisic acid and strigolactones, as well as other important signaling and regulatory molecules (Sierra et al. 2022; Beltran and Wurtzel 2025; Li et al. 2025). In addition, carotenoids are key metabolites that provide provitamin A and dietary antioxidants for humans (Eggersdorfer and Wyss 2018; Rodriguez-Concepcion et al. 2018). The pivotal roles of carotenoids in plants and humans have stimulated intensive research toward understanding the regulatory control of carotenoid accumulation in plants (Cazzonelli and Pogson 2010; Nisar et al. 2015; Rodriguez-Concepcion et al. 2018; Stanley and Yuan 2019; Sun et al. 2022a; Li et al. 2025).

Considerable effort is focused on uncovering genes and factors that control carotenoid accumulation in plants (Stanley and Yuan 2019; Sun and Li 2020; Liang and Li 2023). Phytoene synthase (PSY) catalyzes the first committed step in the carotenoid biosynthesis pathway. PSY serves as a central flux controlling enzyme and its activity strongly influences carotenoid content in crops (Zhou et al. 2022). For example, replacing a daffodil *PSY* with a maize gene encoding a highly active PSY boosts β-carotene levels in Golden Rice by over 20-fold (Paine et al. 2005). Because of its central role, PSY is subject to complex and multilayered regulation in plants (Toledo-Ortiz et al. 2010; Martel et al. 2011; Toledo-Ortiz et al. 2014; Zhou et al. 2015; Álvarez et al. 2016; Lu et al. 2018; Welsch et al. 2018; Rao et al. 2024; Sun et al. 2025).

While a suite of transcription factors and regulators has been shown to control expression of major carotenoid metabolism pathway genes and enzymes (Sun and Li 2020; Liang and Li 2023), other factors are found to contribute to carotenoid accumulation through distinct mechanisms of promoting chromoplast development and enhancing carotenoid sequestration within plastids (Lu et al. 2006; Ariizumi et al. 2014; Tzuri et al. 2015; Chayut et al. 2017; Watkins et al. 2019; Llorente et al. 2020; Stanley et al. 2020; Morelli et al. 2023; Zhou et al. 2023). Despite significant progress, much remains to be uncovered for the genes and factors critical for carotenoid accumulation in plants.

Epistatic interactions of two or more major loci are well-established drivers of fruit flesh colors and root hues in crops. For example, melon (*Cucumis melo* L.) fruit flesh color is primarily controlled by the *Green flesh* (*Gf*) and *White flesh* (*Wf*) loci, with the dominant *Gf* allele epistatic to *Wf* and conferring orange flesh, while *Wf* determines white or green flesh (Hughes 1948; Clayberg 1992). *Gf* encodes CmOr, which regulates chromoplast biogenesis and carotenoid biosynthesis (Lu et al. 2006; Tzuri et al. 2015; Chayut et al. 2017), while *Wf* corresponds to CmRPGE1, a Repressor of Photosynthetic Genes regulating chlorophyll accumulation (Valverde et al. 2025). Three loci (*C1, C2*, and *Y*) interact to determine pepper (*Capsicum annuum* L.) fruit colors from white to red (Hurtado-Hernandez and Smith 1985). *C1, C2,* and *Y* encode Arabidopsis Pseudo-Response Regulator2 (APRR2), PSY1, and capsanthin-capsorubin synthase, respectively (Jeong et al., 2020). In carrot (*Daucus carota* L.), epistasis of two classical loci, *Y* and *Y2,* governs root color variation from white to orange (Iorizzo et al. 2016). Recently, *Y* was found to encode DcRPGE, which functions as a negative regulator of carotenoid biosynthesis by interacting with DcAPRR2 transcription factor (Wang et al. 2024). Members of APRR2 family are known to affect chlorophyll and carotenoid pigmentation colors in fruits including those of *Cucurbitaceae* (Oren et al. 2019; Gebretsadik et al. 2024; Fan et al. 2025; Liu et al. 2025). While individual causal genes at some loci contributing to color epistatic effects have been identified, how they interact and function together in controlling carotenoid accumulation at molecular level remains largely unknown.

*Cucurbita pepo* L (*Cucurbitaceae*) is considered the most economically important species of squash. It encompasses acorn, cocozelle, crookneck, scallop, straightneck, vegetable marrow, and zucchini squash, as well as various pumpkins and ornamental gourds (Paris 1986). Fruit rind and flesh pigmentation are of special interest in *C. pepo.* While rind color is important for the decorative qualities of fall pumpkins and gourds, flesh pigmentation is crucial for quality and nutrition in squash, which can be exceptionally intense in some varieties. Candidates for some loci affecting rind and stem pigmentation have been recently identified (Zhu et al. 2022; Niu et al. 2023; Ding et al. 2024; Gebretsadik et al. 2024). Yet, despite having been described for almost 100 years *in C. pepo*, no major genes responsible for flesh pigmentation have been cloned.

In *C. pepo*, the dominant *Bicolor* (*B*) gene confers precocious yellow fruit coloration (Shifriss 1981). A dominant allele at the *light coloration-2* (*L-2)* locus enhances rind coloration (Paris and Nerson 1986). *B* and *L-2* interact epistatically, resulting in intensely orange-colored fruit rind and flesh and concomitant increase in flesh carotenoids (Schaffer and Boyer 1984; Paris 1988). This interaction has been exploited in the development of varieties with deep orange flesh, such as ‘Orangetti’ (Paris 1993). Notably, *L-2* is part of an allelic series that not only influences pigment intensity, but also exterior patterning such as the reverse striping of ‘Delicata’ (Paris 2020).

In this study, we identified the *B* and *L-2* genes and uncovered the mechanism underlying the synergistic epistasis between *B* and *L-2*, bridging the knowledge gap of epistatic interactions in governing fruit color and carotenoid accumulation in crops.

## Results

### *B* and *L-2* exert an epistatic effect to promote carotenoid accumulation in fruit

Four near-isogenic lines homozygous for the combinations of *B* and *b*, and *L-2* and *l-2*, with genotypes *B/B L-2/L-2*, *B/B l-2/l-2*, *b/b L-2/L-2*, and *b/b l-2/l-2*, have been prepared (Tadmor et al. 2005). Of these four lines, only *B/B L-2/L-2* plants produced mature fruit with deep orange flesh color at 50 days after pollination (DAP). The other three genotypes had light yellow flesh, with the *b/b l-2/l-2* line having the palest yellow flesh color (Fig. 1A).

**Figure 1.**
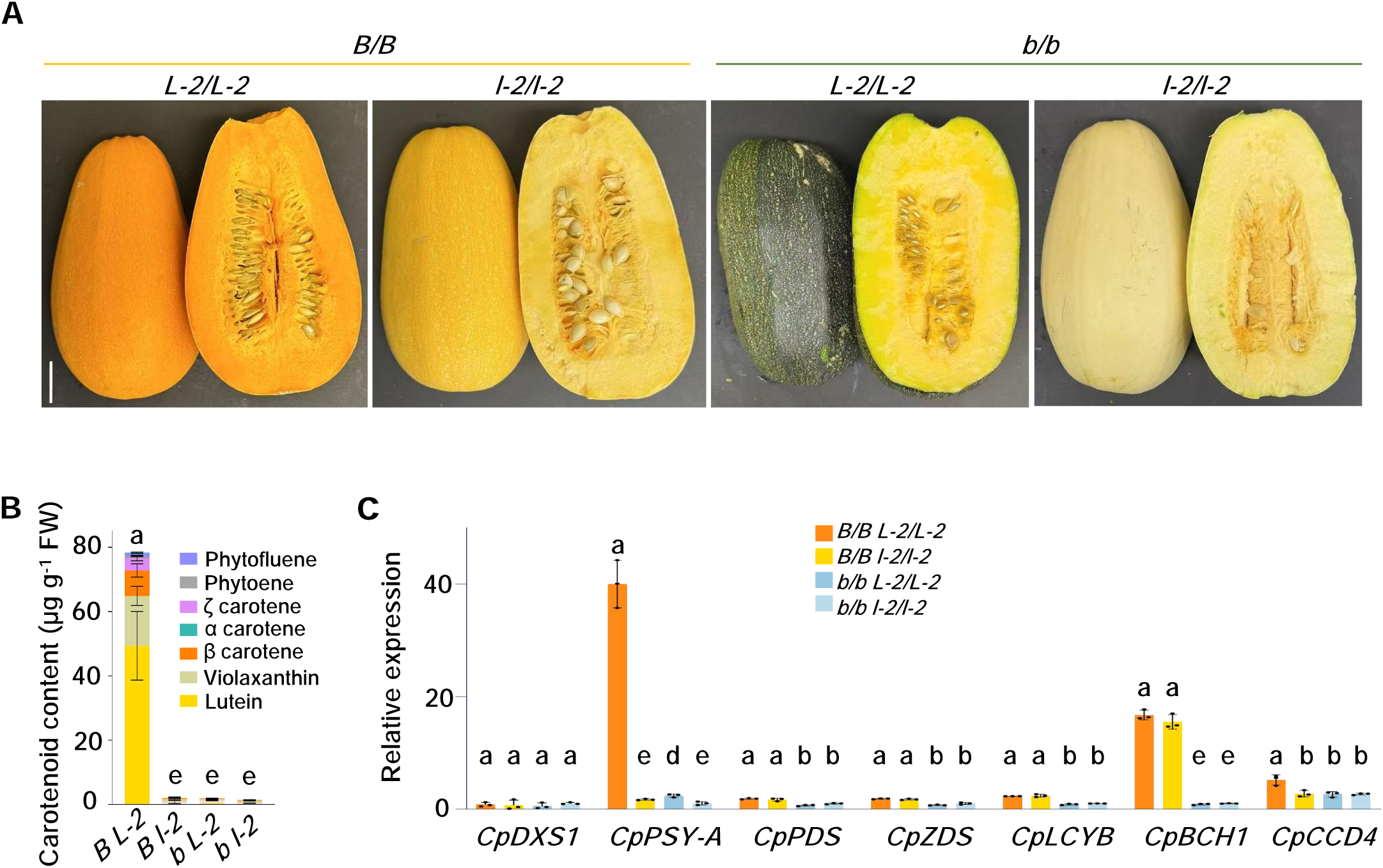
Epistasis of *B* and *L-2* greatly promotes carotenoid accumulation in squash fruit. **A**) Representative fruit phenotype of *B/B L-2/L-2, B/B l-2/l-2, b/b L-2/L-2* and *b/b l-2/l-2* at mature stage (50 days after pollination). Scale bar = 5 cm applicable to all images. **B**) Carotenoid level and composition in mature fruit flesh. *B L-2, B/B L-2/L-2; B l-2, B/B l-2/l-2; b L-2, b/b L-2/L-2; b l-2, b/b l-2/l-2*; FW, fresh weight. **C**) RT-qPCR analysis of expression of carotenoid biosynthesis pathway genes in flesh tissue of mature fruit. *CpDXS1* (1-deoxy-D-xylulose-5-phosphate synthase 1), *CpPSY-A* (phytoene synthase A), *CpPDS* (phytoene desaturase), *CpZDS* (ζ-carotene desaturase), *CpLCYB* (lycopene β-cyclase), *CpBCH1* (β-carotene hydroxylase 1), *CpCCD4* (carotenoid cleavage dioxygenase 4). Data in (**B**) and (**C**) are from three biological replicates and represent as mean ± SD (n = 3). Different letters above bars represent significant differences determined by one-way ANOVA followed by Tukey’s multiple comparison test (*p* < 0.05).

The carotenoid levels and composition in the mature fruit flesh of these four genotypes were analyzed. In the absence of the dominant *L-2,* the *B/B l-2/l-2* mature fruit produced low levels of carotenoids. Similarly in the absence of dominant *B,* the *b/b L-2/L-2* fruit contained a low abundance of carotenoids. However, when *B* and *L-2* were together, an over 35-fold increase of total carotenoids was detected, showing synergistic epistasis effect in stimulating carotenoid accumulation (Fig. 1B and Supplementary Table S1). While lutein was the main carotenoid, an elevated level of β-carotene, the provitamin A carotenoid, was also accumulated in the *B/B L-2/L-2* fruit. The effects of different genotypes on fruit color or carotenoid levels were also clearly observed at 15 DAP stage (Supplementary Fig. S1A). The *B/B L-2/L-2* fruit contained significantly higher total carotenoids than other genotypes already at the fruit enlargement stage and continued to the ripe stage (Supplementary Fig. S1B).

Given the variation in carotenoid content among the genotypes, we analyzed the expression of key carotenoid pathway genes in flesh of mature fruit. *CpPSY-A*, the fruit specific *PSY* gene, showed greatly increased expression in *B/B L-2/L-2* fruit versus other genotypes (Fig. 1C). Several other genes, including *β-carotene hydroxylase 1* (*CpBCH1)*, showed expression differences in the *B/B* versus *b/b* genotypes (Fig. 1C). Among the genes analyzed, *CpPSY-A* was the gene that strongly correlated with total carotenoid content, highlighting its key role in carotenoid production in *C. pepo* fruit.

### *B* encodes a truncated H subunit of Mg chelatase (CpCHLH^B^)

To identify the *B* gene in *C. pepo*, we performed a genotype-by-sequencing analysis of a biparental *C. pepo* F_2_ population segregating for the *B* locus and a population of cultivars that contrast for precocious yellow fruit (Hernandez 2019). SNPs were called from raw data, and an association analysis was conducted. The resulting Manhattan plot localized the *B* gene to the distal end of chromosome 10 (Fig. 2A). The candidate interval for the *B* locus was defined based on the distribution of FDR-corrected association *P*-values along chromosome 10. This region contains 22 annotated genes (Supplementary Table S2, Supplementary Fig. S2). One candidate gene was *Cp4.1LG10g11560*, which encodes the H subunit of magnesium chelatase (CpCHLH). Since the *B* gene causes a precocious yellow phenotype characterized by defective chlorophyll production in ovaries and young fruit (Shifriss 1981), and CpCHLH is involved in chlorophyll biosynthesis, it represents the most plausible candidate. This identification agrees with results from our Israeli co-authors, who mapped *B* using different populations and examined its role in pre-anthesis ovary pigmentation in a related manuscript.

**Figure 2.**
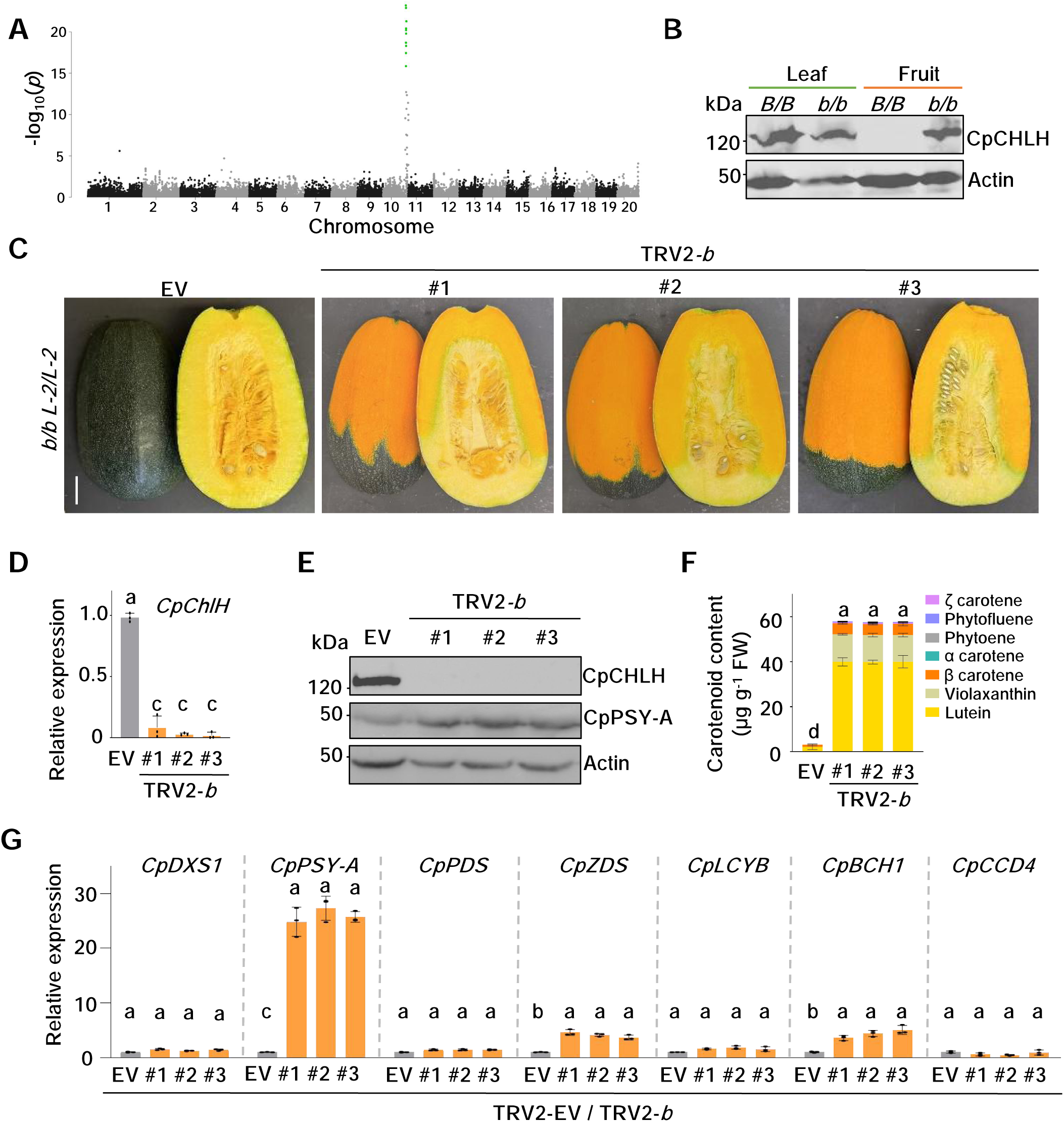
Isolation and characterization of the *B* gene. **A**) Manhattan plot for the *B* locus in *Cucurbita pepo*. The *x*-axis shows genomic positions of SNPs ordered by chromosome, and the *y*-axis shows the -log10(*P*-value) from the association test. A cluster of eleven FDR-significant SNPs on Chromosome 10 is highlighted in green, representing a prominent association peak corresponding to the candidate interval. **B**) Western blot analysis of CpCHLH protein levels in 4-week-old leaves and mature fruit tissue of *B/B L-2/L-2* (*B/B*) and *b/b L-2/L-2* (*b/b*) background. Actin shows protein loading. kDa, protein molecular weight in kilodaltons. **C**) Fruit phenotype of TRV2-*b* in *b/b L-2/L-2* background at 50 DAP stage. EV, empty vector control. #1-#3, three independent virus-induced gene silencing (VIGS) lines. Scale bar = 5 cm applicable to all images. **D**) Transcript levels of *CpChlH* in flesh tissue of EV and TRV2-*b* silenced fruit (orange part) at 50 DAP stage by RT-qPCR. **E**) Western blot analysis of CpCHLH and CpPSY-A protein levels in flesh tissue of EV and TRV2-*b* silenced fruit (orange part) at 50 DAP stage. **F**) Carotenoid level and composition in flesh tissue of EV and TRV2-*b* silenced fruit (orange part). **G**) Relative expression of carotenoids biosynthesis pathway genes by RT-qPCR in flesh tissue of EV and TRV2-*b* silenced fruit (orange part). Data in (**D**), (**F**), and (**G**) are from three biological replicates and represent as mean ± SD (*n* = 3). Different letters above bars represent significant differences determined by one-way ANOVA followed by Tukey’s multiple comparison test (*P* < 0.05).

*B* exhibits tissue-specific effects, impacting fruit but not leaf tissue in *C. pepo* under normal growth conditions (Shifriss 1981). Thus, the full-length coding sequence (CDS) of *CpChlH* from the cDNA libraries of fruit and leaves of *B/B L-2/L-2* and *b/b L-2/L-2* genotypes were amplified and the clones were sequenced via (https://plasmidsaurus.com/). *CpChlH* from *B/B L-2/L-2* leaf tissue showed an identical CDS sequence to that from *b/b L-2/L-2* leaf or fruit. However, we observed a retained 57 bp intron sequence in *CpChlH* from *B/B L-2/L-2* fruit, which was designated as *CpChlH^B^*. The inclusion of the intron sequence after second exon induced a premature stop codon and was predicted to produce a truncated CpCHLH^B^ protein. To confirm this and associate CpCHLH protein accumulation with the *B/B* phenotype, CpCHLH protein abundances from leaf and fruit tissues of the *B/B L-2/L-2* and *b/b L-2/L-2* genotypes were examined by immunoblotting. CpCHLH was detected in leaves of both *B/B* and *b/b* genotypes as well as in *b/b* fruit but undetectable in *B/B* fruit flesh (Fig. 2B), consistent with the *B* gene phenotype of lacking chlorophylls in fruit.

While transformation of *C. pepo* has been reported, the process is still regarded as technically challenging and difficult to carry out successfully. In order to functionally verify the identity of the *B* gene, we performed virus-induced gene silencing (VIGS) of *CpChlH* in the *b/b L-2/L-2* genotype using *Agrobacterium*-mediated cotyledonary injection (Rhee et al. 2022). The *CpChlH* VIGS lines (TRV2-*b*) exhibited a clear phenotypic change, with fruit rind color from green in the empty vector (EV) control to predominantly dark orange and fruit flesh color from light-yellow to dark orange in large sections of the mature fruit (Fig. 2C). Interestingly, a bicolor phenotype was observed in the TRV2-*b* lines with small sectors of fruit that showed the same fruit rind and flesh color as the EV control. That silencing *b* created the bicolor phenotype might be due to reduced *b* dosage as *B/b* heterozygotes have been classically reported to produce bicolor fruit (Shifriss and Paris 1981).

These TRV2-*b* lines were confirmed to be positive with substantially reduced *CpChlH* transcripts and protein levels in the orange flesh parts of the fruit compared to EV controls (Fig. 2D and E). Total carotenoid contents in these tissues were significantly higher from fruit enlargement stage at 15 DAP to mature stage at 50 DAP than EV controls (Fig. 2F; Supplementary Fig. S3A-B). The non-silenced green parts showed similar *CpChlH* transcripts and protein levels as well as total carotenoid content as EV controls (Supplementary Fig. S4A-C).

To investigate how carotenoid biosynthesis pathway genes respond to *CpChlH* silencing, we analyzed their expression in flesh of mature fruit. Among the genes examined, *CpPSY-A* was greatly upregulated in the *CpChlH* silenced fruit tissues in comparison to EV controls, suggesting that silencing *CpChlH* in the *b/b L-2/L-2* background could directly or indirectly affect *CpPSY-A* expression (Fig. 2G). In contrast, upregulation of *CpPSY-A* was not observed in the unsilenced parts of these TRV2-*b* fruit (Supplementary Fig. S4D).

Considering *CpChlH* is a pathway gene for chlorophyll biosynthesis, we investigated whether other genes involved in chlorophyll biosynthesis can also produce the TRV2-*b*-like fruit phenotype. We silenced *glutamyl-tRNA reductase* (*CpGluTR*), *chlorophyllide a oxygenase* (*CpCAO*), *magnesium-chelatase subunit I* (*CpChlI),* and *Genomes Uncoupled 4 (CpGUN4)* in the *b/b L-2/L-2* line via VIGS. Unlike what was observed for TRV2-*b* fruit at an early developmental stage, no changes in fruit phenotypes of these VIGS lines were observed despite their transcript levels being greatly reduced (Supplementary Fig. S5A-J), showing the specific contribution of *CpChlH^B^* to the *B* gene phenotype in *C. pepo* fruit. These findings confirm that *B* encodes CpChlH^B^.

### *L-2* encodes an APRR2 transcription factor

*L-2* synergistically interacts with *B* to enhance carotenoid levels in *C. pepo* fruit. To isolate the *L-2* gene, QTL-seq analysis was utilized to identify the genomic region harboring *L-2* locus. The near-isogenic counterpart F_1_ hybrid ‘Orangetti’ (genotype *B/B L-2/l-2*) was self-pollinated. The F_2_ population of 194 individuals was grown in the field and visually scored for fruit flesh color that was validated by carotenoid measurement. Two bulks with extreme fruit color phenotypes were created by selecting 21 *B/B l-2/l-2* plants that produce pale yellow flesh fruit and 25 *B/B L-2/L-2* individuals that give dark orange fruit. Whole genome resequencing of genomic DNA from the bulks along with the F_1_ parent was carried out and a single QTL for the *L-2* locus was identified, which was located on chromosome 5 (Fig. 3A).

**Figure 3.**
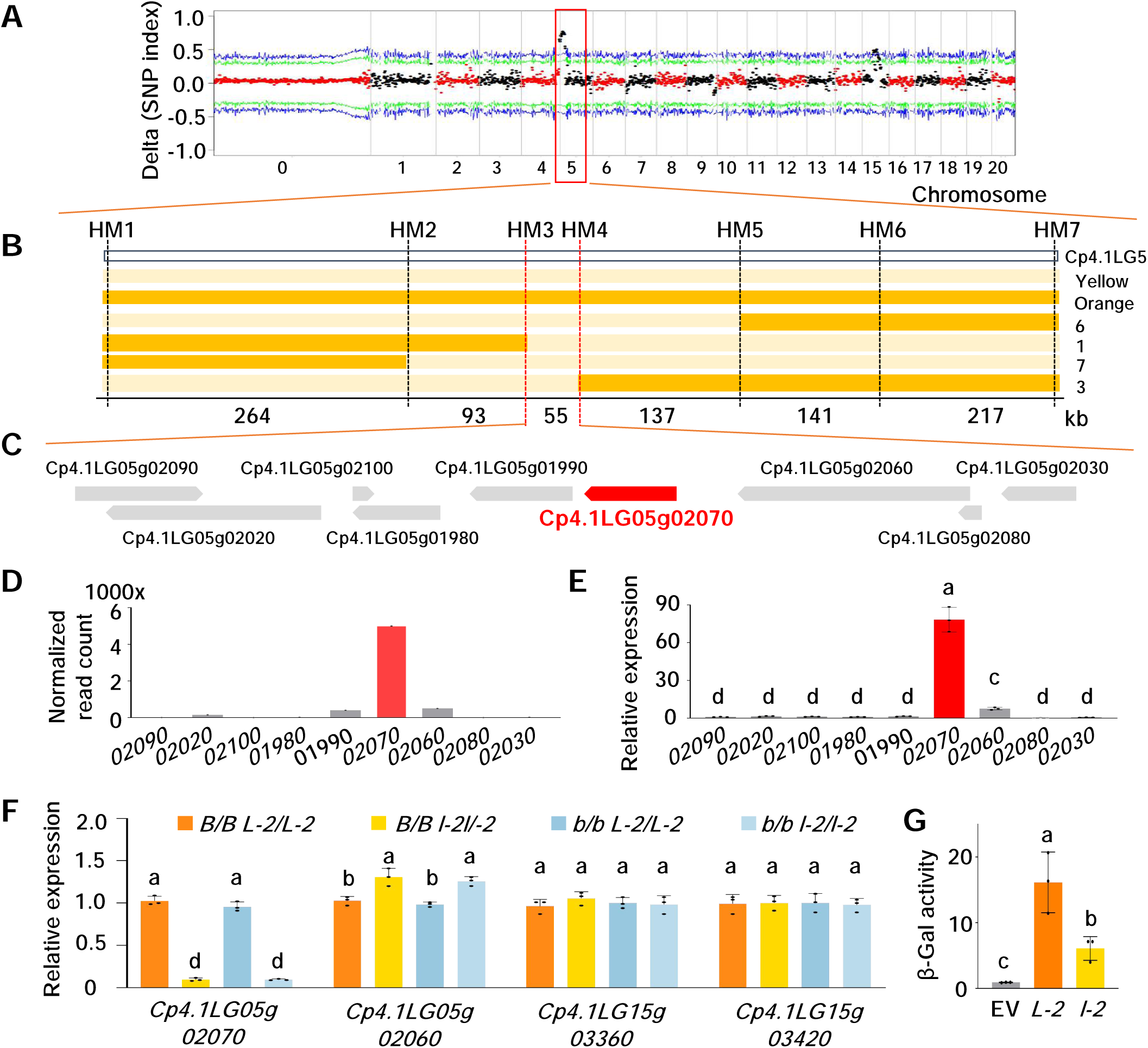
Identification of *L-2* by QTL-seq. **A**) Delta SNP-index plot across *C. pepo* genome. The *x*-axis corresponds to the chromosomal position. Lines are average values of delta SNP-index drawn by sliding window analysis. The upper and lower 99% and 95% confidence thresholds under the null hypothesis of no QTL are indicated by blue and green lines, respectively. Genomic regions where the sliding-window statistic exceeds these thresholds are considered significantly associated with the trait. **B**) Fine mapping of *L-2* on chromosome 5 using indel markers with 26 recombinants. *L-2* was narrowed down within a 55 kb fragment containing 9 genes between markers HM3 and HM4. Dark orange and light-yellow bars represented homozygous chromosomal fragments of orange and yellow genotypes. The 6/1/7/3 on the right is the ID number of recombinant lines that defined the genomic region. Positions of indel markers on the chromosome and physical distances between two markers are indicated. **C**) Nine candidate genes for *L-2* locus. Genes are shown as arrows, with arrow length indicating gene length and arrow direction indicating transcriptional orientation. **D**) Normalized read count of candidate genes in Cpepo_LG at 40 days after pollination were retrieved from the PRJNA339848 dataset (http://cucurbitgenomics.org/). All candidate gene IDs with Cp4.1LG05g that is omitted here. **E**) RT-qPCR confirmation for the expression of all candidate genes in flesh tissue of 50 DAP fruit. **F**) Relative expression of *CpAPRR2* subfamily genes in *C. pepo* analyzed by RT-qPCR. **G**) Transactivation activity of *L-2* and *l-2* was analyzed by β-galactosidase (β-Gal) activity assay. Data in (**E**), (**F**), and (**G**) are from three biological replicates and presented as mean ± SD (n = 3). Different letters above the bars indicate statistically significant differences determined by one-way ANOVA followed by Tukey’s multiple-comparison test (*P* < 0.05).

To map the *L-2* locus, 7 indel markers distributed within the QTL interval were developed and utilized to genotype the 21 F_2_ individuals used to construct the *B/B l-2/l-2* QTL-seq bulk. All the markers were linked to the *L-2* locus. Two flanking markers (HM1 and HM7) were then used to screen approximately 300 F_2_ individuals, of which 26 recombinants were identified. Further genetic mapping with additional five indel markers delineated the *L-2* locus to a 55-kb interval between HM3 and HM4 markers (Fig. 3B).

This 55-kb genomic region contains nine annotated genes with three encoding APRR2 (Fig. 3C, Supplementary Table S3). To identify the candidate gene, we examined the expression profiles of these nine genes from the Cucurbit Genomics Database (http://cucurbitgenomics.org/) (Yu et al. 2023). Only three genes, *i.e. Cp4.1LG05g01990, Cp4.1LG05g02060* and *Cp4.1LG05g02070,* exhibited expression in fruit with *Cp4.1LG05g02070* having the highest expression level (Fig. 3D). The fruit expression patterns of these nine genes were further validated by RT-qPCR (Fig. 3E). *Cp4.1LG05g02070* was annotated to encode an APRR2 protein.

The squash genome contains four members of the APRR2 subfamily. Phylogenetic analysis revealed that they clustered together (Supplementary Fig. S6A). All members displayed a high degree of protein sequence conservation (Supplementary Fig. S6B). Analysis of their expression in fruit tissues of the *B/B L-2/L-2*, *B/B l-2/l-2*, *b/b L-2/L-2*, and *b/b l-2/l-2* near-isogenic lines revealed that *Cp4.1LG05g02070* transcript level was dramatically reduced in *l-2/l-2* as compared with *L-2/L-2*, while the other family genes were similar (Fig. 3F). Based on its fruit-specific expression pattern and differential expression among the four lines, *Cp4.1LG05g02070* that codes for a *CpAPRR2* was selected as the most promising candidate for *L-2,* and named as *CpAPRR2-A*.

To investigate what distinguishes the *L-2* and *l-2* alleles, *CpAPRR2-A* CDS from fruit tissues of the *B/B L-2/L-2* and *B/B l-2/l-2* lines were cloned and sequenced. While *L-2* encoded a full length CpAPRR2-A*, l-2* harbored a 171-bp deletion in the sixth exon of *CpAPRR2-A*, producing a mutated CpAPRR2-A protein with a 57 amino acid deletion (Supplementary Fig. S6C). To investigate whether deletion affects the protein function as a transcription factor, we compared transactivation activity in yeast. CpAPRR2-A from the *L-2* allele exhibited significantly higher β-galactosidase activity or transactivation activity than that from *l-2* (Fig. 3G). These results show that *l-2* encodes a mutated CpAPRR2-A protein with significantly reduced gene expression and transactivation activity.

To functionally validate *CpAPRR2-A* as *L-2*, we silenced *CpAPRR2-A* in the *B/B L-2/L-2* genotype by VIGS. Given the high sequence conservation within the subfamily and to specifically silence the *CpAPRR2-A* gene, we designed the VIGS construct targeting its 3’ untranslated region (UTR). Interestingly, fruit of the *CpAPRR2-A* VIGS (TRV2-*L-2*) lines also exhibited a bicolor phenotype (Fig. 4A). While the unsilenced orange sectors of the TRV2-*L-2* fruit had similar levels of *CpAPRR2-A* expression and carotenoid content (Supplementary Fig. S7A-B), the silenced parts of the TRV2-*L-2* fruit had fruit color changed to green (Fig. 4A), with a greatly reduced *CpAPRR2-A* transcript abundance (Fig. 4B). They also had a significant reduction in total carotenoid levels at the mature stages (Fig. 4C). The TRV2-*L-2* lines also showed a fruit color change and carotenoid level differences at 15 DAP stage (Supplementary Fig. S8A-B). The markedly reduced carotenoid levels in the light-green flesh of TRV2-*L-2* mature fruit verified *CpAPRR2-A* as *L-2* involved in carotenoid accumulation.

**Figure 4.**
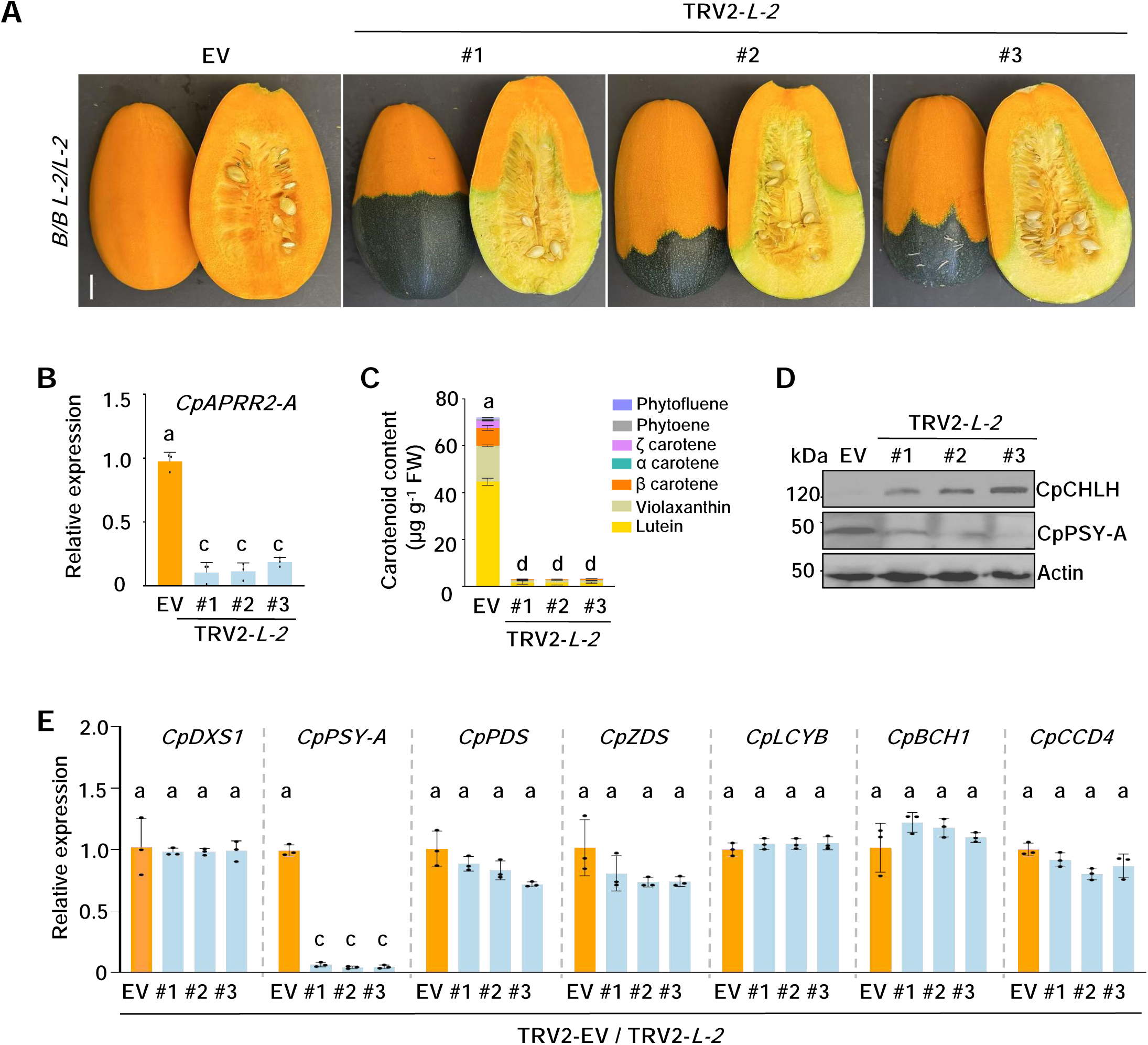
Verification of *L-2* identity by VIGS silencing. **A**) Fruit phenotype of TRV2-*L-2* in *B/B L-2/L-2* background at 50 DAP stage. EV, empty vector control. #1-#3, three independent virus-induced gene silencing (VIGS) lines. Scale bar = 5 cm applicable to all images. **B**) Transcript levels of *CpAPRR2-A* in flesh tissue of EV and TRV2-*L-2* silenced fruit (green part) at 50 DAP by RT-qPCR. **C**) Carotenoid level and composition in flesh tissue of EV and TRV2-*L-2* silenced fruit (green part). **D**) Western blot analysis of CpCHLH and CpPSY-A protein levels in flesh tissue of EV and TRV2-*L-2* silenced fruit (green part). Actin served as a loading control. **E**) Relative expression of carotenoid biosynthesis pathway genes in flesh tissue of EV and TRV2-*L-2* silenced fruit (green part) by RT-qPCR. Data in (**B**), (**C**), and (**E**) are from three biological replicates and presented as mean ± SD (*n* = 3). Different letters above the bars indicate statistically significant differences determined by one-way ANOVA followed by Tukey’s multiple-comparison test (*P* < 0.05).

It is intriguing that suppressing *CpAPRR2-A* in *B/B L-2/L-2* genotype produced a partial dark green rind with light-green flesh fruit that reverted the *B* gene phenotype. We examined the CpCHLH protein and found that it was clearly detectable in the light-green flesh of TRV2-*L-2* fruit but absent in orange sectors (Fig. 4D; Supplementary Fig. S7C). The presence of the CpCHLH protein in the silenced fruit suggests a requirement of *L-2* (*CpAPRR2-A*) for maintaining *B*. To test the hypothesis, we also silenced *l-2*, a mutated and partially functional *CpAPRR2-A,* in the *B/B l-2/l-2* genotype by VIGS. Interestingly, silencing *l-2* also showed the formation of a green section with bicolor fruit phenotype (Supplementary Fig. S9A-B), supporting the requirement of a functional or partially functional *CpAPRR2-A* for the maintenance or formation of *B* in squash fruit.

In addition, transcription of carotenoid pathway genes was examined in the TRV2-*L-2* fruit. Silencing of *CpAPRR2-A* was found to greatly reduce *CpPSY-A* expression, but not other genes examined in the green sectors (Fig. 4E), showing a specific control of *CpPSY-A* by *L-2*. No effects on the pathway gene expression were observed in the non-silenced orange part of the fruit compared to EV controls (Supplementary Fig. S7D). Concomitantly, reduced CpPSY-A protein levels were detected in the green flesh tissues of VIGS positive lines (Fig. 4D) whereas no changes in CpPSY-A protein abundance were found in the orange parts (Supplementary Fig. S7C).

### *B* and *L-2* act in concert to modulate PSY expression and activity

*B* encodes a truncated CpCHLH^B^ and *L-2* encodes CpAPRR2-A. They exert synergistic effects to greatly promote carotenoid accumulation in *C. pepo* fruit. Notably, *CpPSY-A* was expressed highly in the *B/B L-2/L-2* line (Fig. 1C), whereas its expression was greatly increased in TRV2-*b* silenced lines (Fig. 2G) and reduced in TRV2-*L-2* plants (Fig. 4E), showing that epistatic interaction between *B* and *L-2* regulated *CpPSY-A* expression. To further investigate the mechanism underlying the synergistic action of *B* and *L-2*, we examined CpPSY-A protein levels in fruit flesh tissues of the *B/B L-2/L-2* and *b/b L-2/L-2* lines. A higher abundance of CpPSY-A protein was detected in *B/B L-2/L-2* compared to *b/b L-2/L-2* at both 15 and 50 DAP stages (Fig. 5A), indicating that *B* promotes CpPSY-A protein accumulation in the *L-2* background.

**Figure 5.**
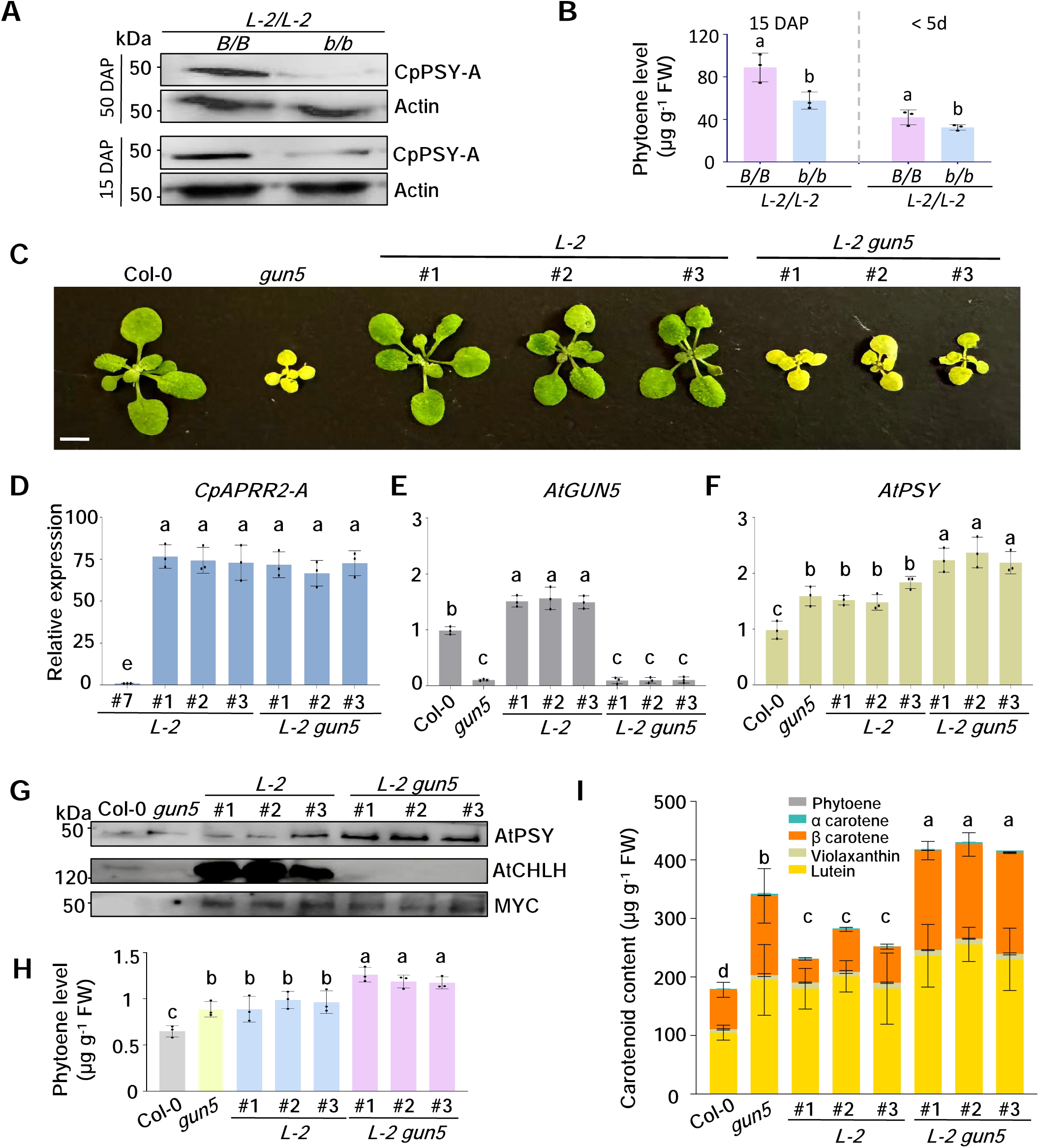
*B* (*gun5*) and *L-2* affect *PSY* expression. **A**) Western blot analysis of CpPSY-A protein levels in *B/B L-2/L-2* and *b/b L-2/L-2* squash fruit at 15 and 50 DAP stages. Actin served as a loading control. **B**) PSY activity in squash fruit measured after NFZ (norflurazon) treatment. The *B/B L-2/L-2* and *b/b L-2/L-2* fruit at young (< 5 d) and 15 DAP stages were treated with 10 μM NFZ overnight. Phytoene levels quantified by UCP^2^ are used as a proxy for PSY enzymatic activity. **C**) Phenotype of Arabidopsis wild type Col-0, *gun5* mutant, and three independent *L-2* overexpression transgenic lines in Col-0 (*L-2* #1-3) or *gun5* (*L-2 gun5* #1-3) background. Scale bar = 2 cm applicable to all images. **D-F**) Relative expression of *CpAPRR2-A*, *AtGUN5* and *AtPSY*, respectively, in Col-0, *gun5*, *L-2*, *L-2 gun5* lines. Gene expression is relative to the lowest expressing transgenic line (*L-2* #7 in **D**) or Col-0 (**E** and **F**), which was set to 1. **G**) PSY and CHLH protein levels in Col-0, *gun5*, L-2, and *L-2 gun5* lines. MYC was used as a reference to normalize loading of L-2 protein level. **H**) NFZ treatment on Col-0, *gun5*, *L-2*, *L-2 gun5* lines to detect PSY activity changes. **I**) Total carotenoids extracted from Col-0, *gun5*, *L-2*, *L-2 gun5* lines and measured by UPC^2^. Data in (**D**-**F**), (**H**), and (**I**) are from three biological replicates and presented as mean ± SD (*n* = 3). Letters above bars indicate significant differences determined by one-way ANOVA followed by Tukey’s multiple-comparison test (*P* < 0.05).

To investigate whether the increased PSY protein level at 15 DAP leads to enhanced enzymatic activity, we treated the young fruit of the *B/B L-2/L-2* and *b/b L-2/L-2* lines with norflurazon (NFZ), an assay commonly used to monitor *in vivo* PSY activity and metabolic flux through the carotenoid biosynthesis pathway (Rodríguez-Villalón et al. 2009; Zhou et al. 2015). NFZ is an inhibitor of phytoene desaturase (PDS) and blocks phytoene conversion into downstream metabolites, leading to phytoene accumulation. The NFZ-treated fruit accumulated significantly higher levels of phytoene in *B/B L-2/L-2* than *b/b L-2/L-2* during early fruit development, which was observable at extremely young stage (Fig. 5B). The result supports that the increased PSY protein is enzymatically active.

To examine whether the interaction between an impaired *ChlH* and *APRR2* leads to increased *PSY* expression and protein accumulation across plant species, we also overexpressed *L-2* (*CpAPRR2-A*) in Arabidopsis wild-type Col-0 and in a *gun5* mutant background. The *gun5* mutant carries a defect in *AtChlH* (Strand et al. 2003), thus genetically mimicking *C. pepo B*. Three T_3_ homozygous lines for *L-2* and three for *L-2 gun5* were selected (Fig. 5C). *L-2* was similarly overexpressed among these selected transgenic lines in both wild-type and *gun5* background (Fig. 5D). *AtChlH* was remained at low levels in the transgenic lines with *gun5* background (Fig. 5E). Notably, *AtPSY* transcript level was enhanced more than 2-fold in the *L-2 gun5* lines, and also significantly higher in the *gun5* mutant with an endogenous *APRR2* gene (Fig. 5F). A high abundance of PSY protein was detected in *L-2 gun5* lines compared to Col-0, *gun5,* and *L-2* (Fig. 5G). NFZ treatment of leaf tissues revealed that PSY activities were coordinately increased in these lines (Fig. 5H). Moreover, higher total carotenoid levels were observed in the *gun5* mutant and *L-2* lines, with a further increase detected in the *L-2 gun5* transgenic lines (Fig. 5I). These data corroborate the findings in squash that interaction between an impaired *ChlH* and *APRR2* upregulates *PSY* expression and activity to promote carotenoid accumulation across plant species.

### Only CpCHLH^b^, not CpCHLH^B^, physically interacts with CpAPRR2-A

To investigate how *B* and *L-2* together enhance CpPSY-A expression and activity, protein-DNA and protein-protein interaction experiments were performed. A previous study reported that carrot DcAPRR2 directly binds to the promoter of *DcPSY* to activate its transcription (Wang et al. 2024). We performed yeast one-hybrid (Y1H) and dual-luciferase (pDual-LUC) assays to examine the activation of the *CpPSY-A* promoter by CpAPRR2-A. Yeast growth on selective medium was observed when *CpPSY-A* promoter and *CpAPRR2-A* (*L-2*) constructs were co-transformed, whereas no growth was observed with controls (Fig. 6A). In addition, the *CpPSY-A* promoter-driven luciferase showed strong signals when co-transformed with *CpAPRR2-A* in comparison with EV and a pathway gene (*CpDXS1)* as controls (Fig. 6B). The *in vivo* transactivation activity was quantified with dual luciferase reporters of nanoLUC and fireflyLUC (Sun et al. 2025), and the *CpPSY-A* promoter showed significantly higher activation by *CpAPRR2-A* than with EV or *CpDXS1* (Fig. 6C). These results showed that CpAPRR2-A was directly bound to the *CpPSY-A* promoter and transactivated its expression. We also found that the *l-2* encoded protein retained binding activity to the *CpPSY-A* promoter, but its transactivation of the *CpPSY-A* promoter was greatly reduced compared with the *L-2* encoded one (Supplementary Fig. S10A-D). These results support the ability of CpAPRR2-A in activating *CpPSY-A* expression, consistent with that reported in carrot (Wang et al. 2024).

**Figure 6.**
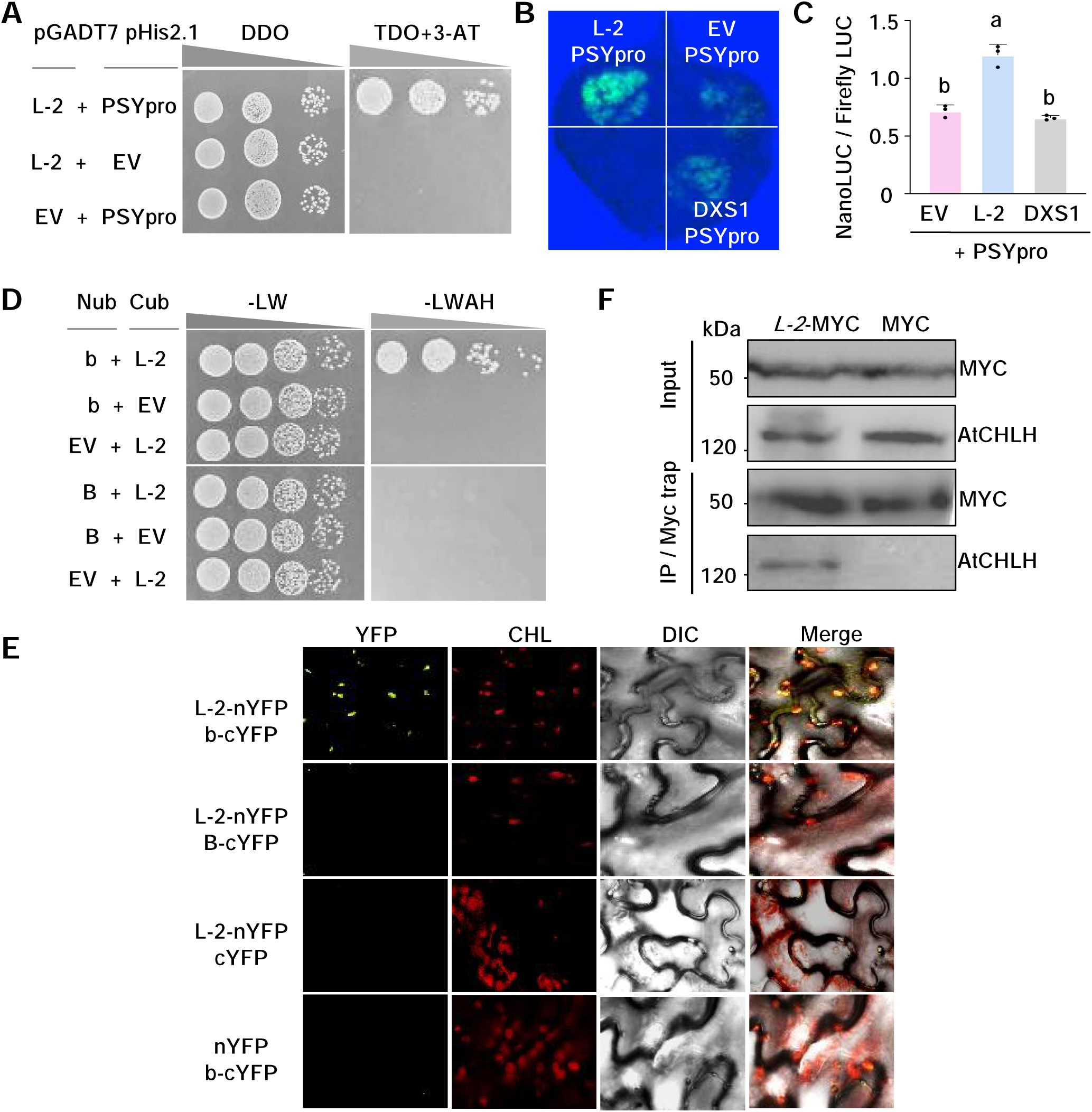
CpAPRR2-A activates *CpPSY-A* expression and physically interacts with CpCHLH^b^ but not CpCHLH^B^. **A**) Y1H (yeast one-hybrid) analysis showed the binding of CpAPRR2-A (L-2) to *CpPSY-A* promoter (PSYpro) *in vitro*. Yeast cells were co-transformed with the PSYpro-pHis2.1 construct and L-2-pGADT7 plasmid, spread on DDO (double dropout medium, SD/–Leu/–Trp) and TDO (triple dropout medium, SD/–Leu/–Trp/–His with 80 mM 3-AT) for selection. Empty-vector (EV) controls show no growth. **B**) Transactivation assay in *N. benthamiana* leaves showing *L-2* activation of *CpPSY-A* promoter. EV and an unrelated protein CpDXS1 were used as negative controls. **C**) Luciferase activity was quantified using protein extracts treated with luciferin or coelenterazine. Transactivation activity was normalized using FireflyLUC and expressed as NanoLUC/FireflyLUC with three biological replicates. Different letters above the bars indicate statistically significant differences (one-way ANOVA followed by Tukey’s multiple comparison test, *P* < 0.05). **D**) Y2H (yeast two-hybrid) assay of CpCHLH^b^ (b)/ CpCHLH^B^ (B) interactions with CpAPRR2-A (L-2). B or b was fused to the N-terminal half of ubiquitin (Nub), while L-2 was fused to the C-terminal half (Cub). Yeast co-transformants were spotted in 10-fold serial dilutions onto nonselective (-LW) plate to confirm plasmid maintenance and selective plate (-LWAH) to assess interaction, followed by photograph after 3 days. **E**) BiFC analysis of b or B interactions with L-2. L-2 was fused to nYFP, and b or B was fused to cYFP. L-2-nYFP/cYFP and nYFP/b-cYFP served as negative controls. Reconstituted YFP signals were detected by confocal microscopy. Scale bar = 20 µm applicable to all images. **F**) Co-IP (co-immunoprecipitation) assay of b interaction with L-2. Total proteins were extracted from 3-week-old Arabidopsis leaves expressing Myc-tagged *L-2* in Col-0 background or a Myc-tagged unrelated protein (negative control). Extracts were immunoprecipitated with anti-Myc beads and analyzed by immunoblotting with anti-Myc and anti-CHLH antibodies on both input and IP samples.

CpPSY-A gene expression and protein levels were highly abundant in the *B/B L-2/L2* line and substantially low in the *b/b L-2/L-2* line (Fig. 1 and Fig. 5A). To examine whether *B* and *b* differentially affect the capacity of CpAPRR2-A in regulating CpPSY-A gene expression and protein levels, protein-protein interactions of their encoded CpCHLH^B^ and CpCHLH^b^ with CpAPRR2-A were examined. When CpAPRR2-A (L-2) and CpCHLH^b^ (b) were co-expressed in yeast, the yeast grew well on the selective medium. However, when CpAPRR2-A and CpCHLH^B^ (B) were expressed together in yeast, no yeast growth was observed on the selective medium (Fig. 6D). These results revealed that the CpAPRR2-A physically interacted with CpCHLH^b^, but not with CpCHLH^B^ in yeast.

To confirm this specific interaction *in vivo*, a bimolecular fluorescence complementation (BiFC) assay was carried out in *Nicotiana benthamiana* leaves. Yellow fluorescent signals were clearly observed when CpCHLH^b^ and CpAPRR2-A were co-expressed, while no signals were detected when CpCHLH^B^ and CpAPRR2-A were co-expressed (Fig. 6E), further supporting that CpCHLH^b^ but not CpCHLH^B^ physically interacted with CpAPRR2-A. The fluorescent signals colocalized with chlorophyll, indicating that the interaction occurs in plastids, the site for carotenoid biosynthesis (Sun et al. 2019).

To further verify the interaction between CpCHLH^b^ and CpAPRR2-A *in planta*, a coimmunoprecipitation (co-IP) assay was performed using Myc-tagged proteins from Arabidopsis lines expressing *L-2* in Col-0 background and an unrelated gene generated previously in our lab as a negative control (Rao et al. 2024). With similar amounts of loading and eluted Myc fusion proteins, the CHLH protein was coimmunoprecipitated by CpAPRR2-A from protein extracts of the *L-2* line but not detected in the negative control (Fig. 6F), supporting the interactions between CpCHLH^b^ and CpAPRR2-A *in planta*.

### CpCHLH^b^ suppresses the ability of CpAPRR2-A in activating *CpPSY-A* expression

To investigate whether the differential interactions of CpAPRR2-A with CpCHLH^b^ and CpCHLH^B^ play a role in affecting the ability of CpAPRR2-A to regulate *CpPSY-A* expression, we conducted a competitive binding assay with and without CpCHLH^b^ or CpCHLH^B^. The *CpPSY-A* promoter-driven luciferase exhibited strong signals when the promoter construct was co-expressed with *L-2* in comparison with EV control. No increased signals were observed when the promoter construct was co-transformed with *b* or *B* construct *in planta* (Fig. 7A), indicating no transactivation of *CpPSY-A* by CpCHLH^b^ or CpCHLH^B^. When the *CpPSY-A* promoter was co-expressed with *L-2* and *b* constructs, luciferase signal was markedly reduced compared to co-expression with *L-2* and *B* constructs (Fig. 7A). The *in vivo* transactivation activity was quantified using the NanoLUC/FireflyLUC dual-luciferase system by normalizing nanoLUC with the firefly LUC activity. Consistently, the results showed that CpCHLH^b^ significantly reduced the transactivation activity of CpAPRR2-A on the *CpPSY-A* promoter in comparison with CpCHLH^B^ (L-2+b vs L-2+B; Fig. 7B). These results indicate that CpCHLH^b^ and CpAPRR2-A interaction suppresses the activity of CpAPRR2-A in activating *CpPSY-A* expression.

**Figure 7.**
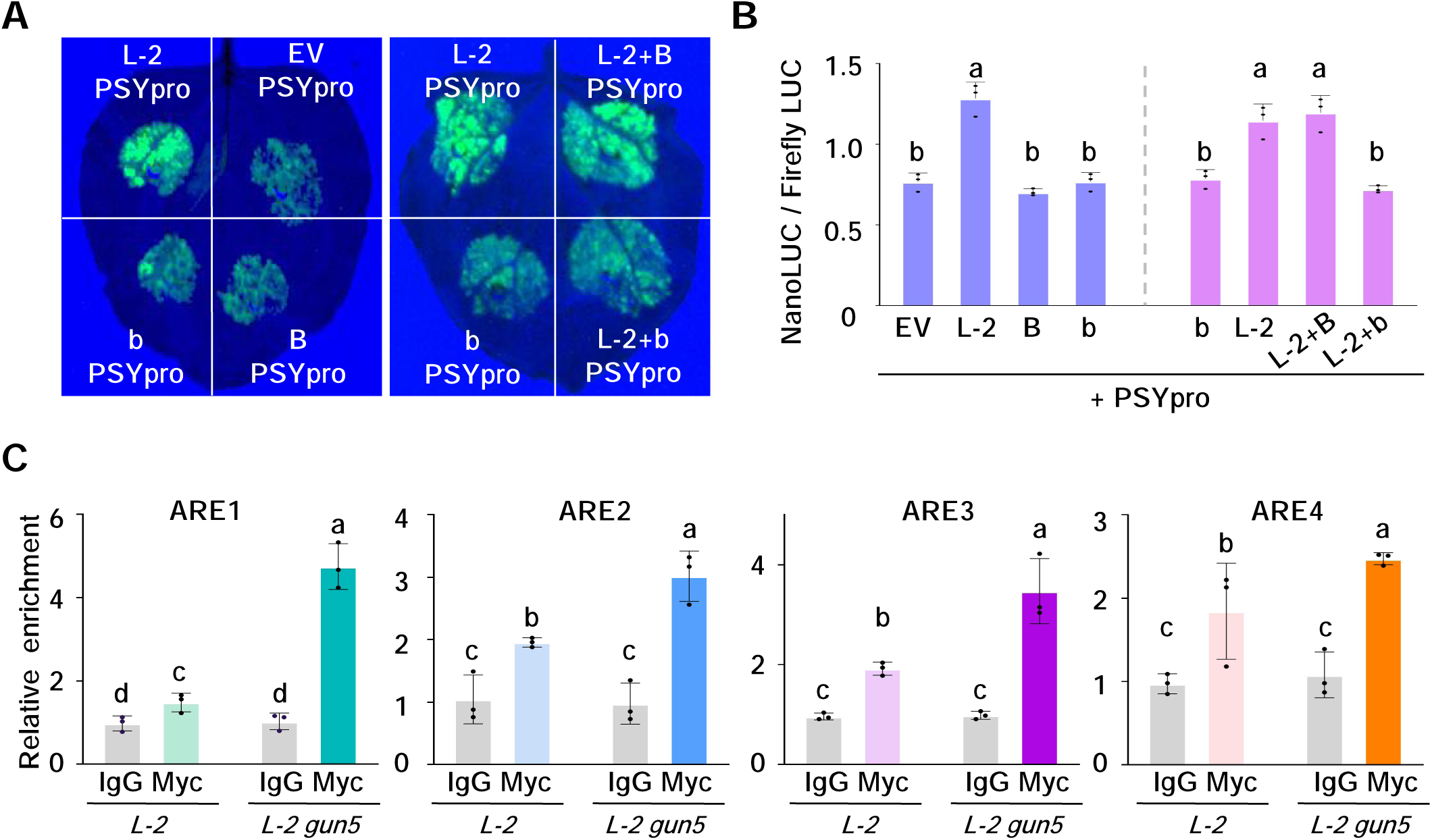
CpCHLH^b^ but not CpCHLH^B^ suppresses CpAPRR2-A-induced *CpPSY-A* expression. **A**) Transactivation assay in *Nicotiana benthamiana* leaves. Agrobacterium cells carrying the CpAPRR2-A promoter (PSYpro)-LUC reporter were co-infiltrated with effectors expressing CpAPRR2-A (L-2), CpCHLH^b^ (b), CpCHLH^B^ (B), or the L-2+B and L-2+b combinations. Empty vector (EV) was used as a negative control. LUC signal was detected 48 h after infiltration, and representative leaf images are shown. **B**) Quantification of transactivation activity using the NanoLUC/FireflyLUC dual-luciferase system. Luciferase activity was measured from protein extracts treated with luciferin for NanoLUC or coelenterazine for FireflyLUC. **C**) ChIP-qPCR analysis of L-2 binding to the *PSY* promoter in Col-0 and *gun5* backgrounds. Chromatins from *L-2* and *L-2 gun5* plants were immunoprecipitated with anti-Myc antibody or IgG (control). Enrichment of *PSY* promoter fragments was quantified by qPCR and expressed as 1.5-2% input DNA. Data in (**B**) and (**C**) are from three biological replicates and presented as mean ± SD (*n* = 3). Different letters denote significant differences determined by one-way ANOVA followed by Tukey’s multiple-comparison test (*P* < 0.05).

To corroborate these findings, chromatin immunoprecipitation followed by qPCR (ChIP-qPCR) was carried out. Chromatins from Arabidopsis transgenic *L-2* and *L-2 gun5* lines were immunoprecipitated with anti-Myc antibody or IgG (control). The binding peaks of CpAPRR2-A to ARE1-ARE3 (APRR2-recognition element 1-3) motifs of the Arabidopsis *PSY* promoter were markedly lower with chromatins from the *L-2* line than the *L-2 gun5* line (Fig. 7C). The results indicate that functional CHLH in the *L-2* line suppresses CpAPRR2-A binding to the *PSY* promoter, whereas the mutated CHLH in *L-2 gun5* promotes CpAPRR2-A binding to the promoter. These findings imply that while the interaction of CpAPRR2-A with CpCHLH^b^ prevents its binding to the promoter of *CpPSY-A*, the lack of interaction with CpCHLH^B^ protein reduces the inhibitory effect, allowing CpAPRR2-A to activate *CpPSY-A* expression for carotenoid accumulation.

## Discussion

The intricate epistatic interactions among major genetic loci are fundamentally important in shaping the color variations observed in fruits and roots. Major genetic loci underlying squash flesh pigmentation have been recognized for nearly a century (Hernandez 2019), yet their molecular identities and mechanisms remain large unknown. Here, we reveal the identities of two classical color loci, *B* and *L-2*, and elucidate the mechanism of their epistatic interaction in producing intensely orange-fleshed fruit with high levels of carotenoid accumulation in *C. pepo*. The *B* gene encodes a truncated Mg-chelatase subunit H, CpCHLH^B^, and *L-2* encodes transcription factor CpAPRR2-A. The *b* allele encodes functional CpCHLH^b^ protein that interacts physically with CpAPRR2-A to suppress its activation of *CpPSY-A* and inhibits carotenoid biosynthesis. The epistasis of *B* and *L-2* arises from lack of interaction between truncated CpCHLH^B^ protein and CpAPRR2-A, thereby enabling CpAPRR2-A to activate *CpPSY-A* expression. Subsequently, high *CpPSY-A* expression leads to greatly enhanced carotenoid production to produce intensely orange fleshed fruit. Mechanistic models among various genotypes in regulating carotenoid accumulation with different flesh color phenotypes are shown in Figure 8. These findings not only provide insight into epistatic mechanisms governing fruit carotenoid accumulation but also offer a promising molecular framework for future breeding strategies aimed at improving the nutritional value and visual quality of food crops.

**Figure 8.**
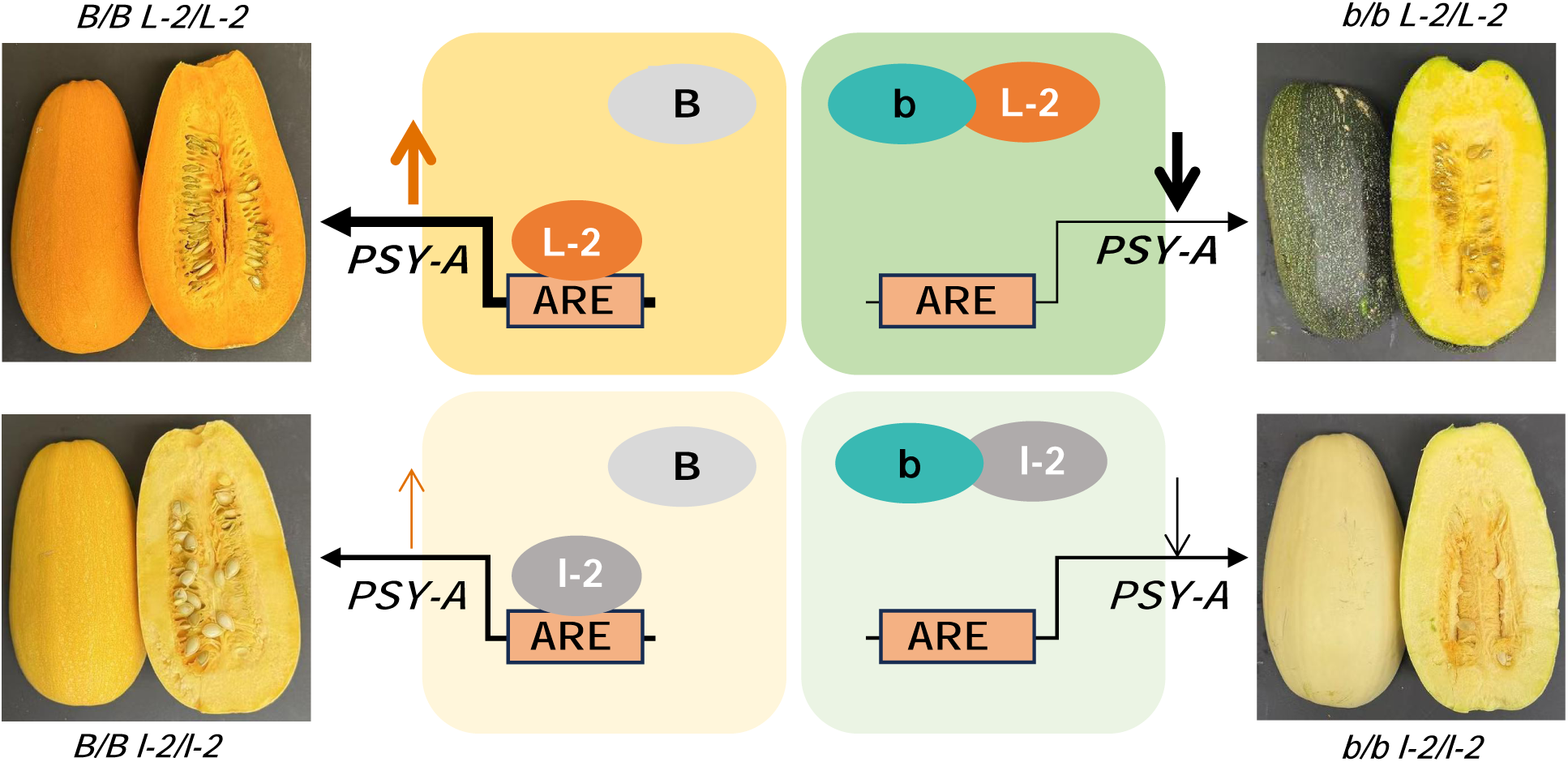
Mechanistic model of *B* and *L-2* epistasis in coordinating *CpPSY-A* regulation to drive carotenoid production in squash fruit. In *B/B L-2/L-2* genotype, the dominant *B* encoded protein CpCHLH^B^ (B) loses interaction with CpAPRR2-A (L-2), allowing it to freely bind to the *CpPSY-A* promoter to activate *CpPSY-A* transcription and promote carotenoid accumulation to develop intense orange fruit. In contrast, in the *b/b L-2/L-2* genotype, the recessive *b* encoded CpCHLH^b^ (b) interacts with CpAPRR2-A and prevents its activation of *CpPSY-A*, resulting in green rind fruit with low carotenoids. In *B/B l-2/l-2* and *b/b l-2/l-2* genotypes, while the *l-2* encoded protein retains partial activity and can still interact with CpCHLH^b^, its low gene expression and low transactivation activity makes it indistinguishable in affecting *CpPSY-A* expression in *B/B l-2/l-2* and *b/b l-2/l-2* genotypes, producing pale-yellow fruit phenotypes with low carotenoid content.

### The dominant *B* allele produces a truncated CpCHLH^B^

*B* can exert profound effects to increase flesh carotenoid levels in *C. pepo* (Shifriss 1981, Schaffer et al., 1984). The dominant *B* allele produces truncated CpCHLH^B^ with undetectable CpCHLH protein in the fruit (Fig. 2). Since CHLH is essential for chlorophyll biosynthesis as the catalytic and substrate-binding subunit of magnesium chelatase, it is not surprising that *B* leads to the precocious yellow fruit phenotype, which is described in detail in a related manuscript. However, the loss of chlorophyll accumulation alone cannot explain the subsequent increase in carotenoid accumulation in the developing fruit flesh, revealing the importance of interaction of *B* with *L-2*.

VIGS is an effective tool to verify gene functions in cucurbits (Rhee et al. 2022). Interestingly, silencing of *CpChlH* in the *b/b L-2/L-2* genotype produced bicolor fruit phenotype. The bicolor phenotype may result from reduced *b* dosage as *B/b* heterozygotes are classically reported to produce bicolor fruit (Shifriss and Paris 1981). *ChlH* silencing or mutation typically induces pale-yellow phenotypes in leaves due to impaired chlorophyll biosynthesis (Hiriart et al. 2002; Zhang et al. 2024). However, silencing *CpChlH* here did not visibly affect leaf pigmentation but caused a pronounced color change phenotype in fruit (Fig. 2C), suggesting variable effects of *CpChlH* in chlorophyll biosynthesis in different tissues of *C. pepo*. A possible explanation is differential sensitivity to the reduced CpCHLH activity in regulating chlorophyll biosynthesis in leaf as compared with fruit tissue. The *B* gene conferred fruit phenotype appears specific to *CpCHLH^B^*, as silencing other chlorophyll biosynthesis pathway genes did not produce the same fruit phenotype.

### *L-2* encodes an APRR2 transcription factor regulating pigment production in fruit

*L-2* is the synergistic partner of *B* to produce intense orange fruit flesh with high carotenoid accumulation in *C. pepo*. *L-2* was identified as *Cp4.1LG05g02070* that codes *CpAPRR2-A* with fruit-specific high expression. Interestingly, silencing *CpAPRR2-A* in both *B/B L-2/L-2* and *B/B l-2/l-2* genotypes resulted in chlorophyll biosynthesis, reverting CpCHLH^B^ into CpCHLH^b^, suggesting that the formation of *B* requires a functional or partially functional CpAPRR2-A although the molecular basis remains to be elucidated. In addition, silencing also produced a bicolor fruit phenotype, unlike *APRR2* silencing that produces pale-yellow or light-green fruit in tomato (Chen et al. 2025) and pepper (Jeong et al. 2020). While how this happens remains to be elucidated, the bicolor phenotype suggests a balance and interaction between *B* and *L-2* in governing fruit color in *C. pepo*.

APRR2 homologs are broadly conserved across plant species, related to but distinct from the Golden2-like transcription factors (GLK2) that modulate pigment biosynthesis and chloroplast development (Waters et al. 2009; Sun et al. 2025). APRR2 impacts fruit color in many crops, including fruit rind and/or flesh color in several cucurbits, such as cucumber (*Cucumis sativus* L.) (Fan et al. 2025; Liu et al. 2025), melon and watermelon (Oren et al. 2019; Yue et al. 2025), and *C. pepo* (Gebretsadik et al. 2024). While APRR2 homologs have been shown to affect chlorophyll and carotenoid pigments in fruit flesh and peel, the mechanism underlying their regulatory role remains largely unknown. A recent study in carrot revealed that DcAPRR2 exerts its impact by directly interacting with the promoters of *DcPSY* to control carotenoid biosynthesis (Wang et al. 2024). We also confirmed that CpAPRR2-A bound to the promoter of *CpPSY-A* to activate its transcription, which underscores the important role of APRR2 in regulating *PSY* expression for carotenoid production.

### Epistasis of *B* and *L-2* results from loss of interaction between their encoded proteins, allowing CpAPRR2-A to activate *CpPSY-A* expression

While epistasis is a widespread feature controlling fruit and root pigmentation in diverse crop species, the mechanistic insights remain largely unknown. In tomato, the epistasis of *tangerine* (*t*) over *yellow-flesh* (*r*) was suggested to involve *cis*-carotenoids in a feedback regulation of *SlPSY1* to affect fruit color (Kachanovsky et al. 2012). Notably, *CpPSY-A* was expressed highly in *B/B L-2/L-2* fruit and substantially upregulated in TVR2-*b* fruit in *b/b L-2/L-2* genotype (Figs 1C, 2G), showing that *B* affects CpAPRR2-regulated *CpPSY-A* expression. This effect is proved to be conserved across species, as Arabidopsis plants with a combination of *gun5* and *APRR2* similarly increased *AtPSY* expression (Fig. 5F), reinforcing a conserved mechanism.

The mechanism underlying such regulation stems from the physical interaction of CpCHLH^b^ with CpAPRR2-A which prevents CpAPRR2-A from binding to the promoter of *CpPSY-A* for activating its expression. However, truncated CpCHLH^B^ disrupts this interaction, enabling CpAPRR2-A to strongly activate *CpPSY-A* for elevated carotenoid production (Fig. 8). This mechanism underpins the epistatic control of *B* and *L-2* in producing intensely orange-fleshed fruit in squash. In carrot, DcRPGE1 interacts with DcAPRR2 to similarly suppress DcAPRR2-mediated transcriptional activation of the *DcPSY* gene, whereas mutated DcRPGE1 fails to bind DcAPRR2 resulting in carotenoid biosynthesis (Wang et al. 2024). While *DcRPGE* was identified as the *Y* locus, the identity of *Y2* remains unknown for the epistatic interaction of *Y* and *Y2* in controlling carrot root colors.

The molecular basis of *B* and *L-2* epistasis in boosting carotenoid production involves CpCHLH^B^ and CpAPRR2-A interplay in regulating *CpPSY-A* expression. On the other hand, the mutated CpCHLH^B^ may also affect plastid fate to contribute to high levels of carotenoid accumulation in squash. CHLH not only serves as a core component in chlorophyll biosynthesis but also, as genomes uncoupled 5 (GUN5), fulfills a key role in plastid-to-nucleus retrograde signaling to affect chloroplast development (Mochizuki et al. 2001; Strand et al. 2003). The *gun5* mutation impairs chloroplast development (Mochizuki et al. 2001; Strand et al. 2003). CpCHLH^B^ in *B* fruit with a defect in CpCHLH also attenuates chlorophyll biosynthesis and chloroplast development with precocious yellow fruit coloration. The weakened chloroplast development in *B* fruit cells may be prime plastids for carotenoid accumulation when pathway activity is upregulated during fruit ripening. Concurrently, *B* also enables CpAPRR2-A to upregulate *CpPSY-A* expression and metabolic flux into carotenoid synthesis, further promoting carotenoid accumulation in chromoplasts, the plastids specialized for storing carotenoids.

Indeed, compromised photosynthetic capacity or weakened chloroplast identity has been shown to change plastid fate and increase chromoplast competency for carotenoid accumulation (Llorente et al. 2020). Such enhanced chromoplast competency would provide a more favorable cellular environment for carotenoid sequestration and stabilization (Egea et al. 2010; Sun et al. 2018; Li et al. 2025; Wrightstone et al. 2025), thereby amplifying pigment accumulation beyond the effect of transcriptional activation alone. The markedly elevated carotenoid accumulation by *B* and *L-2* in squash fruit results from their synergistic action to regulate biosynthetic activity and may be facilitated by potentially *B* altered plastid fate, forming a coordinated mechanism that drives high levels of carotenoid accumulation. This coordinated mechanism could help explain the synergistic action observed from our studies of ORANGE (OR) and PSY for high levels of carotenoid accumulation (Sun et al. 2021). OR conditions chromoplast development, but its impact on total carotenoid content depends on the pathway activity and is greatly augmented by increased PSY activity (Wrightstone et al. 2025).

Collectively, our results elucidate an epistatic gene pair (*B* and *L-2*) that coordinates carotenoid accumulation to control fruit pigmentation in *C. pepo*. As a duo, *B* and *L-2* make squash a very rich source of carotenoids. This work not only uncovers the identities of two classical color loci but also bridges a long-standing gap in understanding the molecular mechanisms underlying epistatic interplay into the coordinated regulation of carotenoid accumulation in crops.

## Methods

### Plant materials and growth conditions

The *Cucurbita pepo* materials used in this study were four nearly isogenic inbreds of genotypes *B/B L-2/L-2*, *B/B l-2/l-2*, *b/b L-2/L-2*, and *b/b l-2/l-2* (Tadmor et al. 2005). These inbreds were in the genetic background of ‘Vegetable Spaghetti’ with genotype *b/b l-2/l-2*, to which *B* and *L-2* were introgressed to the sixth backcross generation, followed by several generations of self-pollination. The donor parent of *B* and *L-2* was a breeding line named Precocious Fordhook Zucchini (Shifriss and Paris 1981). In addition, for QTL-seq analysis, an F_2_ mapping population was generated by self-pollinating the ‘Orangetti’ F_1_ hybrid squash. ‘Orangetti’, which was first commercialized in 1986 (Paris et al. 1985; Paris 1993), is homozygous for *B* and heterozygous for *L-2* (genotype *B/B L-2/l-2*). Both of its parents are nearly isogenic to the four homozygous nearly isogenic lines and were derived in the same fashion, with one being of genotype *B/B L-2/L-2* and the other of genotype *B/B l-2/l-2*. Further, Golden Zucchini and Romulus F2 population and a cultivated diversity panel were used.

The squash plants were grown in Cornell University facilities either at 14 h light / 10 h dark and 26 / 20 °C (day/night) in the Guterman greenhouse facility in Ithaca, New York or in the field at Homer C. Thompson farm in Freeville, New York. All plants were grown according to standard horticultural practices. All fruits were hand pollinated with tagging to record the number of days after pollination (DAP) and collected at 15 DAP (fruit enlargement) and 50 DAP (mature) stages.

*Arabidopsis thaliana* Columbia-0 (Col-0) and *Nicotiana benthamiana* plants were grown in growth chambers maintained at 16 h light / 8 h dark and 23/20 °C (day/night). Arabidopsis *gun5* mutant was obtained from Arabidopsis Biological Resource Center (ABRC; Cat. CS6499). Tissues were collected, frozen immediately in liquid nitrogen, and stored at - 80 °C until use.

For norflurazon (NFZ) treatment, 0.1 g of fresh fruit flesh tissues were finely chopped (< 1 mm pieces) and fully submerged with 10 μM NFZ (Sigma; Cat. 34364) overnight in the dark. For Arabidopsis leaves, approximately 20 fresh leaves from 3-week-old plants were soaked in 70 μM NFZ solution under dark for 2 h, followed by transferring to 10 μM NFZ solution under light for 4 h as described (Zhou et al. 2015).

### Carotenoid analysis

Carotenoids from fruit flesh tissues or NFZ-treated samples were extracted using the method as described previously (Barja et al. 2021) with some modifications. Briefly, 0.2 g fruit samples were extracted in 1 mL of hexane: acetone: methanol (2:1:1, v/v/v), and partitioned with 0.4 mL of ddH_2_O, followed by centrifugation at 12,000 g for 10 min at 4 °C. The upper pigment-containing phase was collected and dried down under vacuum. Carotenoids from 0.15 g Arabidopsis leaf samples were extracted with acetone and ethyl acetate using the method (Sun et al. 2025).

Saponification of esterified carotenoids in fruit samples was carried out as described (Zhang et al. 2014) with slight modifications. The dried extracts were resuspended in 300 μL of 10% (w/v) KOH in methanol and incubated in the dark for 30 min. After adding 400 μL of ddH_2_O, the mixture was vortexed and partitioned with 600 μL of ethyl acetate. Followed by centrifugation at 12,000 g for 10 min at 4 °C. The upper organic phase was extracted again with 600 μL of ethyl acetate and the combined upper phases were evaporated to dryness under vacuum.

Extracted carotenoids were analyzed using a Waters UPC^2^ as detailed (Yazdani et al. 2019; Hazra et al. 2025). The individual carotenoids were identified based on their retention times and their characteristic absorption spectra in comparison with standards and quantified using standard calibration curves.

### RNA isolation and quantitative real-time PCR analysis

Total RNA from squash flesh/leaf or Arabidopsis leaf samples were extracted using TRIzol reagent (Invitrogen; Cat.15596018). RNA was reverse transcribed using an MMLV Reverse Transcriptase, GPR (TaKaRa Bio; Cat. 639576). RT-qPCR was carried out with iTaq^TM^Universal SYBR^®^ Green Master Mix (Bio-Rad; Cat. 1725124) on a CFX384 Touch Real-time PCR Detection System (Bio-Rad) with gene-specific primers (Supplementary Table S4) as detailed (Chayut et al. 2021). Three biological and three technical replicates were tested for each sample.

### Genotyping-by-Sequencing (GBS), SNP calling, and association analysis of *B*

GBS data from an F_2_ population (’Golden Zucchini’ × ‘Romulus’; n = 76) and a cultivated diversity panel (n = 104) were used to map the *B* locus. SNPs were called with the TASSEL-GBS pipeline using enzyme-specific parameters and merged with published *Cucurbita pepo* GBS data (Hernandez et al. 2023). Prior to imputation, SNPs were filtered with a minor allele frequency ≥ 0.05, heterozygosity ≤ 0.9, and missing data ≤ 0.5. Missing genotypes were then imputed using the TASSEL LDKNNiImputationHetV2Plugin. Following imputation, the same SNP filters were reapplied and individuals with more than 20% missing data were removed. GWAS was performed in GAPIT using a mixed linear model with a marker-based kinship matrix (K) and the first two principal components derived from K as covariates. The B locus interval on chromosome 10 was defined from an FDR-corrected cluster of associated SNPs, which served as the basis for subsequent candidate gene analysis.

### Western blot analysis

Total proteins from squash flesh and Arabidopsis leaf samples were extracted as described previously (Hazra et al. 2025). Western blot analysis was carried out as described (Sun et al. 2025). The primary antibodies used included anti-PSY antibody (1:400 dilution) generated by Abgent (San Diego, CA), anti-CHLH (1:1,000 dilution; Agrisera; Cat. AS224712), anti-Myc antibody (1:2,000; VWR; Cat. 103628-444), and anti-Actin (1:1000 dilution; Sigma-Aldrich; Cat. A0480). Protein bands were visualized with the WesternBright ECL kit (Advansta, USA; Cat. K12045D20).

### QTL-seq analysis and genetic mapping of *L-2*

For QTL-seq analysis, an F_2_ mapping population was generated by self-pollinating the ‘Orangetti’ F_1_ hybrid (genotype *B/B L-2/l-2*) and a total of 193 F_2_ individuals were grown in the field. Mature fruit flesh color was visually evaluated and carotenoid content from three fruit per plant was measured by UPC^2^ (Yazdani et al. 2019). Two bulks were created using 21 plants with light yellow flesh and 25 plants with dark orange flesh fruit phenotype. Genomic DNA from ‘Orangetti’ and the two bulks with equal amount of leaf materials from individual plants were extracted using the cetyltrimethylammonium bromide (CTAB) method. QTL-seq library construction and whole genome resequencing with paired-end 2 × 150 bp reads were carried out at Azenta Life Sciences (https://www.azenta.com/). Sequencing read processing, alignment to the reference *C. pepo* genome (Montero-Pau et al. 2018), and QTL identification were carried out as described previously (Tan et al. 2020).

To develop indel markers covering the *L-2* QTL region, unique ∼600-bp flanking genome sequences containing insertion and deletion (indel) polymorphisms with size differences over 20 bp were selected. Primers were synthesized and amplified in the F_1_ parent and the two bulks for polymorphism. The developed indel markers were used to genotype first in the individuals composing of the yellow bulk and then additional ∼ 300 F_2_ plants to identify recombinants, which were grown to fruit maturity. Based on phenotypic and genotypic data and the physical distances among markers, a map was constructed according to their physical positions in the genome. All marker primers are listed in Supplementary Table S4.

### Virus-Induced Gene Silencing (VIGS) analysis

Tobacco rattle virus (TRV)-based VIGS was carried out following the procedure described (Rhee et al. 2022). To silence target genes in *C. pepo*, the 300 bp 3’UTR region of *CpAPRR2-A*, 500-bp CDS of *CpChlH,* and 300-bp CDS of *CpGluTR, CpCAO, CpChlI1*, or *CpGUN4* were cloned into TRV RNA2 vector (Senthil-Kumar and Mysore 2014). The constructs were transformed into *Agrobacterium tumefaciens* strain GV3101 by electroporation. Agrobacterium cultures containing a TRV2 gene construct or empty vector as the negative control were infiltrated with TRV1 (1:1, v/v, OD_600_ = 1) into the cotyledons of the *C. pepo* seedlings (at least 30 plants for each gene construct). Positive lines were confirmed by RT-qPCR with primers binding outside the VIGS target region primers (Supplementary Table S4). Fruit phenotype was examined 3-4 weeks after infiltration.

### Generation of Arabidopsis transgenic lines

To generate *L-2* overexpression lines, the coding sequence (CDS) of *CpAPRR2-A* without a stop codon was amplified using gene-specific primers (Supplementary Table S4) and cloned into pGWB17 vector (Rao et al. 2024). The plasmid was electroporated into *Agrobacterium* GV3101 and transformed into wild type *Arabidopsis thaliana* Col-0 or the *gun5* mutant (CS6499, ABRC) via the floral dip method. Positive transgenic plants were identified by western blot with an anti-Myc antibody (1:2,000 dilution; VWR; Cat. 103628-444). Three independent homozygous T_3_ lines for each background (*L-2* in Col-0 and *L-2* in *gun5*) were selected for further experiments.

### Yeast one hybrid (Y1H) assay

Y1H analysis was carried out as described (Sun et al. 2025). To generate the *CpPSY-A* promoter DNA bait construct, a 2,000-bp fragment upstream of the *CpPSY-A* start codon was amplified and inserted upstream of the HIS3 reporter gene in pHIS2.1 vector (Clontech). To produce pGADT7-*L-2* and pGADT7-*l-2* prey constructs, coding sequences of *L-2* and *l-2* were amplified and fused in-frame with the GAL4 activation domain in pGADT7 (Clontech). Yeast strain AH109 was co-transformed with pairs of plasmids. Transformants were grown on SD/-Leu/-Trp medium and subsequently tested for HIS3 reporter activation on selective medium SD/-Leu/-Trp/-His supplemented with 80 mM 3-amino-1,2,4-triazole (3-AT) at 30°C for 2-3 days.

### Transactivation activity assay

To compare the transactivation activity difference between *L-2* and *l-2*, coding sequences of *L-2* and *l-2* were fused in frame with the GAL4 DNA-binding domain in the pDEST32 vector. The constructs and empty vector were transformed into yeast strain YRG-2, which carries the GAL4pro:LacZ reporter, and selected on SD/−Leu/−Trp dropout medium. To measure β- galactosidase activity, cells from overnight yeast cultures were lysed by freeze-thaw cycles. Their transactivation activity (Miller units) in catalyzing O-nitrophenyl-β-D-galactopyranoside was measured as described (Sun et al. 2025).

### Dual reporter assay

To construct pDual reporter construct, a 2 kb *CpPSY-A* promoter region upstream of the start codon was amplified (Supplementary Table S4), cloned into Gateway pENTRY vector pDONR207, and then to the destination luciferase reporter vector pGWB401-nanoLUC-UBQ10-FLUC as described (Sun et al. 2025). The UBQ10 promoter-driven firefly luciferase served as a stable internal reference for quantification. To generate the effector constructs, coding sequences of *L-2, l-2*, and *DXS1* were cloned into pGWB17. The reporter and effector constructs were electroporated into *Agrobacterium* GV3101 individually, which were mixed in pairs in a 1:1 ratio (OD_600_ = 0.1-0.2), and infiltrated into 3-week-old leaves of *Nicotiana benthamiana* plants. After 2-3 days, the infiltrated leaves were sprayed with 50 μM D-luciferin (ThermoFisher; Cat.88291) or coelenterazine (RPI; Cat. C61500) and nanoLUC or FLUC signals were detected by a ChemiDoc MP system (Bio-Rad). For quantification, bioluminescent signal intensities were measured using ImageJ software and the reporter activation was quantified following normalization by FLUC activity. All experiments were performed with three biological replicates.

### Yeast two-hybrid (Y2H)

As CHLH is a membrane localized protein, a split-ubiquitin system for Y2H assay was performed as previously described (Sun et al. 2022b). The *L-2* coding sequence, lacking the transit peptide and stop codon, was cloned into pMetYCgate to generate the Cub fusion, while *B* and *b* coding sequences from *B/B* and *b/b* lines, respectively, without the transit peptide and stop codon sequences were inserted into pNXgate for Nub fusions. The Cub fusion constructs were transformed into yeast THY.AP5 and pNXgate constructs into THY.AP4. After mating, the resulting diploid yeast cells were plated out on a drop-out medium SD/-Leu/-Trp for growth control and on SD/-Leu/-Trp/-His/-Ade selective medium lacking leucine (L), tryptophan (W), adenine (A), and histidine (H) supplemented with 0.05 mM Met at 30°C for 2-3 days. A β-galactosidase assay using ortho-nitrophenyl D-galactoside (oNPG) as the substrate was carried out to examine interaction strength as detailed (Yuan et al. 2015).

### Co-immunoprecipitation (Co-IP)

Co-IP was carried out as described (Rao et al. 2024). Briefly, total proteins were extracted from leaves of Arabidopsis expressing either L-2-Myc or an unrelated protein-Myc fusion and incubated with anti-Myc magnetic beads (Miltenyi Biotec, Auburn, CA; Cat. 130-091-123). The bead-protein mixtures were captured by μ-columns and eluted out with 2× SDS loading buffer before resolving in SDS-PAGE. Immunoblotting was performed as described above for western analysis.

### Bimolecular Fluorescence Complementation (BiFC)

BiFC analysis was performed as detailed (Sun et al. 2022b). In short, the coding sequences of *L-2* and *b* or *B* were cloned into the Gateway-compatible binary vectors pSITE-nYFP and pSITE-cYFP, respectively, to generate N- or C-terminal YFP fusion constructs. *Agrobacterium* GV3101 cultures carrying the nYFP and cYFP constructs were mixed in pairs at a 1:1 ratio and infiltrated into four-week-old *Nicotiana benthamiana* leaves. YFP fluorescence signals were examined 48 h post-infiltration under a confocal laser scanning microscope (Leica TCS SP5) with YFP excited at 488 nm and emitted between 520-560 nm and chlorophyll autofluorescence in the 620-680 nm range. Images were acquired and processed using Leica LAS AF software, and identical settings were applied to all treatments to allow direct comparison of fluorescence intensity.

### Chromatin Immunoprecipitation followed by quantitative PCR (Chip-qPCR)

ChIP assay was carried out using the EpiQuik Plant ChIP Kit (EpigenTek, Cat. P-2014-24) following the manufacturer’s protocol. Approximately 2 g of finely grounded fresh leaf samples from Arabidopsis wild type and *gun5* mutant overexpressing L-2-Myc fusion protein were cross-linked with 1% formaldehyde under vacuum for 10 min, followed by quenching with 0.125 M glycine. Chromatin released from nuclei was sheared to 200-600 bp fragments on ice using a Bioruptor sonicator (cycles of 40 s ON / 40 s OFF). Equal amounts of chromatin from each sample were incubated overnight at 4°C with either anti-Myc antibody, 1:2,000 dilution; VWR, Cat. 103628-444) for L-2 bound to DNA or anti-H3K4me3 antibody (Millipore, Cat. 07-473) as a positive histone modification control. Antibody-chromatin complexes were immunoprecipitated with protein A/G magnetic beads. Cross-links between proteins and DNA were reversed at 65°C, and the recovered DNA was used for qPCR analysis using primer pairs targeting the ARE1-ARE4 regulatory elements within the *PSY* promoter (Supplementary Table S4). Enrichment levels were calculated relative to input DNA and normalized to a non-binding genomic region.

### Statistical analysis

GraphPad Prism version 8.4 software was used for statistical analysis. Data were analyzed using one-way ANOVA followed by Tukey’s test. Significance of differences are indicated by different letters. All experiments were carried out with at least three biological replicates.

## Accession Numbers

L-2 (Cp4.1LG05g02070); B (Cp4.1LG10g11560); CpDXS1 (Cp4.1LG17g01110); CpPSY-A (Cp4.1LG01g19670); PDS (Cp4.1LG08g06310); ZDS (Cp4.1LG04g01620); LCYB (Cp4.1LG15g07370); BCH1 (Cp4.1LG09g00740); CCD4 (Cp4.1LG01g01120); CHLI (Cp4.1LG19g06700); CHLD (Cp4.1LG16g08680); CHLG (Cp4.1LG10g07890); GUN4 (Cp4.1LG12g05090); GluTR (Cp4.1LG11g05040); CAO (Cp4.1LG13g04640); AtPSY (AT5G17230); AtGUN5 (AT5G13630).

## Supplemental Data

**Supplementary Figure S1**. *B and L-2* together significantly enhance carotenoid accumulation at fruit developmental stage.

**Supplementary Figure S2**. *B* candidate region on chromosome 10.

**Supplementary Figure S3**. TRV2-*b* silencing fruit at fruit developmental stage.

**Supplementary Figure S4**. Analysis of green sections of the TRV2-*b* fruit.

**Supplementary Figure S5**. Silencing of chlorophyll biosynthetic pathway genes by virus-induced gene silencing (VIGS).

**Supplementary Figure S6**. APRR2 subfamily and *CpAPRR2-A* transcript variations from *L-2* and *l-2*.

**Supplementary Figure S7**. Analysis of yellow sections of the TRV2-*L-2* fruit.

**Supplementary Figure S8**. TRV2-*L-2* silencing fruit at fruit developmental stages.

**Supplementary Figure S9**. Silencing of *l-2* in *B/B l-2/l-2* geneotype.

**Supplementary Figure S10**. *l-2* encoded mutated CpAPRR2-A with 57 amino acid deletion has reduced binding and transactivation activity.

**Supplementary Table S1**. Carotenoid level and composition in flesh tissue of fruit at 50 DAP.

**Supplementary Table S2**. Annotated genes in the candidate region of *B* locus.

**Supplementary Table S3**. Annotated genes in the candidate region of *L-2* locus.

**Supplementary Table S4**. Primers used in this study.

## Acknowledgments

We thank Ms. Tara Fish and Dr. Theodore Thannhauser for their assistance with carotenoid analysis, and the Plant Cell Imaging Center (PCIC) at Boyce Thompson Institute for the access of microscopes, as well as the Freeville Farm crew and Guterman greenhouse staff for plant care and technical support. We thank undergraduate Kyle Griswold for assistance with experiments, and additional undergraduate researchers in Mazourek’s lab for their help with the squash saponification experiment.

## Author contributions

L.L and M.M designed the study; L.X and X.Z performed most of the experiments and analyzed the data; E.W and K.M helped with *L-2* gene mapping; G.I, C.H, and A.G managed plant genetic materials and helped with *B* gene identification, Z.F. carried out bioinformatics analysis of QTL-seq data. H.S.P provided genetic materials and guided bulk sample selection; A.S, J.M, and L.C assisted in data analysis and/or revised the manuscript; L.X, L.L and M.M wrote the paper with contributions from all coauthors.

