## Supplementary figures for "Epistasis of two classical color genes, *B* and *L-2*, synergistically controls carotenoid accumulation in squash"

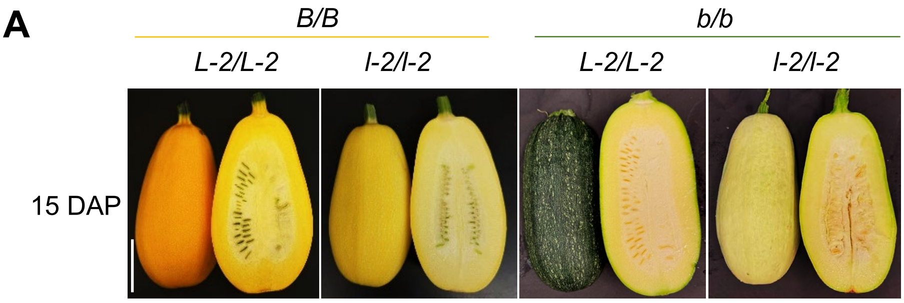


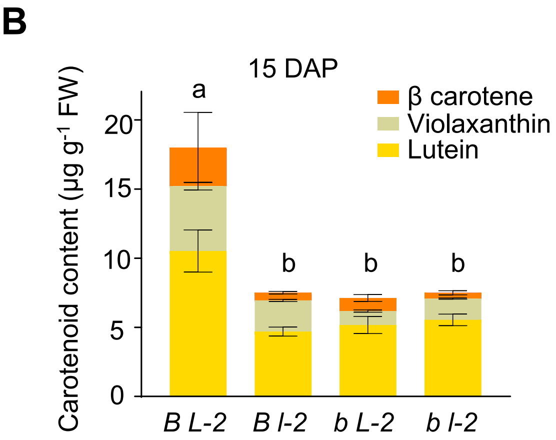


**Supplementary Figure S1.** *B* and *L-2* together significantly enhance carotenoid accumulation at fruit developmental stage. **A)** Representative fruit phenotype of *B/B L-2/L-2* , *B/B l-2/l-2*, *b/b L-2/L-2* and *b/b l-2/l-2* at 15 DAP (days after pollination) stage, scale bar = 5 cm applicable to all images. **B)** Carotenoid content and composition for *B/B L-2/L-2* (*B L-2*), *B/B l-2/l-2* (*B l-2*), *b/b L-2/L-2* (*b L-2*) and *b/b l-2/l-2* (*b l-2*) at 15 DAP stage. Data are from three biological replicates and presented as means ± SD (n = 3). Different letters above the bars indicate statistically significant differences, as determined by one-way ANOVA followed by Tukey's multiple-comparison test ($p < 0.05$).

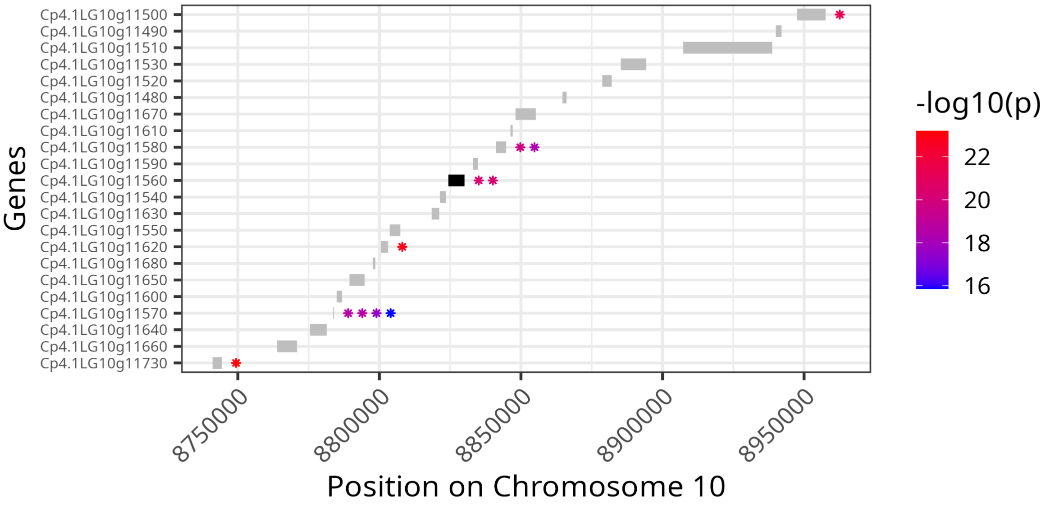


**Supplementary Figure S2.** *B* candidate region on chromosome 10. The eleven FDR-significant SNPs are plotted relative to annotated genes within the ~200 kb candidate region (Chr10: 8.75-8.95 Mb). Stars indicate SNP locations, colored by -log10(*p*) value. The gene model corresponding to gene *Cp4.1LG10g11560* is shown in black. All SNPs fall within annotated genes, consistent with the GBS-derived marker distribution.

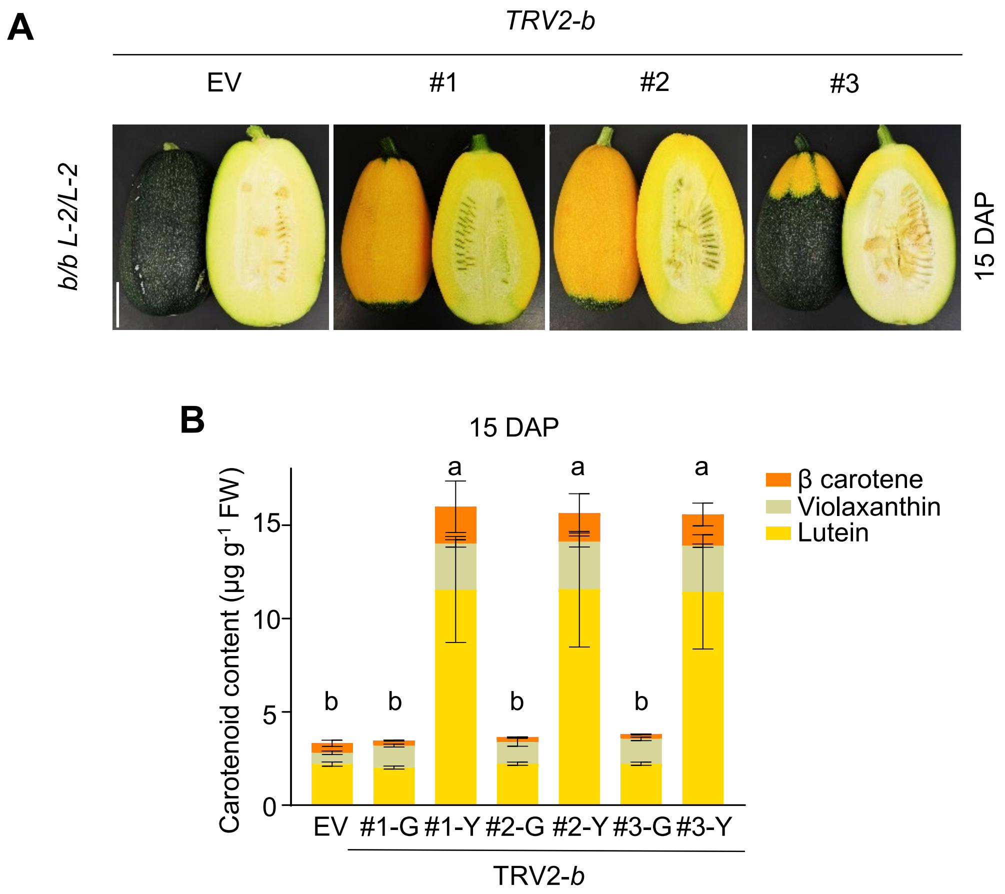


**Supplementary Figure S3.** TRV2-*b* silencing fruit at fruit developmental stage. **A)** Fruit phenotype of TRV2-*b* lines in *b/b L-2/L-2* background at fruit enlargement (15 DAP) stage. EV (Empty Vector) as control, scale bar = 2 cm applicable to all images. **B)** Carotenoid content and composition at 15 DAP stage. G represents the green flesh part of the fruit, and Y represents the yellow part. Data in (**B**) are from three biological replicates and presented as mean ± SD (n = 3). Different letters denote significant differences, as determined by one-way ANOVA followed by Tukey's multiple-comparison test ($p < 0.05$).

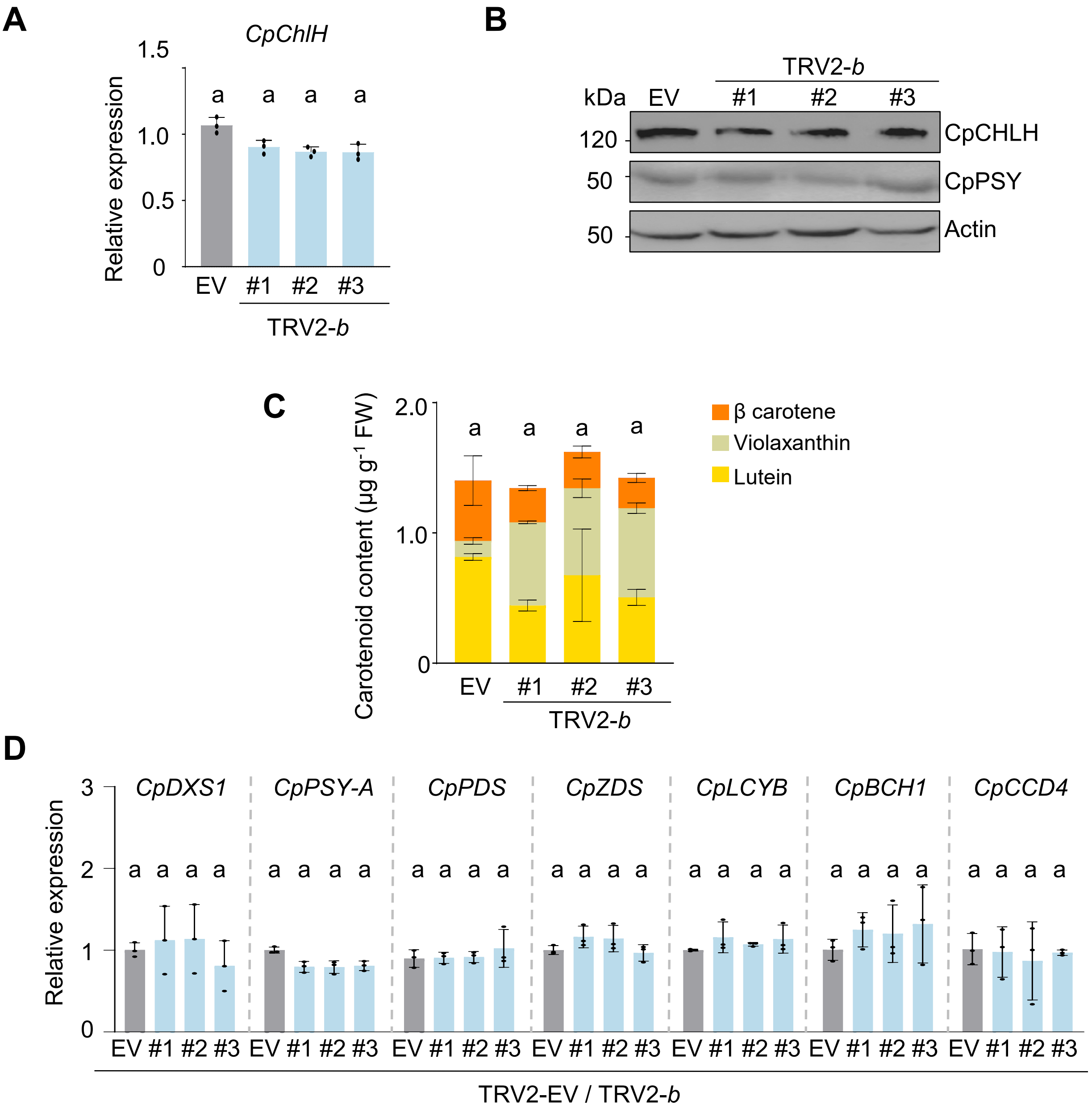


**Supplementary Figure S4.** Analysis of green sections of the TRV2-*b* fruit. **A)** The transcription level of *CpChlH* in green parts of empty vector (EV) control and TRV2-*b* fruit in *b/b L-2/L-2* background. **B)** CpCHLH and CpPSY protein levels were detected by western blot in fruit (green part) of TRV2-*b* lines at 50 DAP stage. Actin served as a loading control. **C)** Carotenoid content and composition in green parts of EV and TRV2-*b* fruit at 50 DAP stage. **D)** Relative expression of carotenoid biosynthesis pathway genes was detected by RT-qPCR in green part of fruit at 50 DAP stage. Data in (**A**), (**C**), and (**D**) are from three biological replicates and presented as mean ± SD (n = 3). Different letters denote significant differences, as determined by one-way ANOVA followed by Tukey's multiple-comparison test ($p < 0.05$).

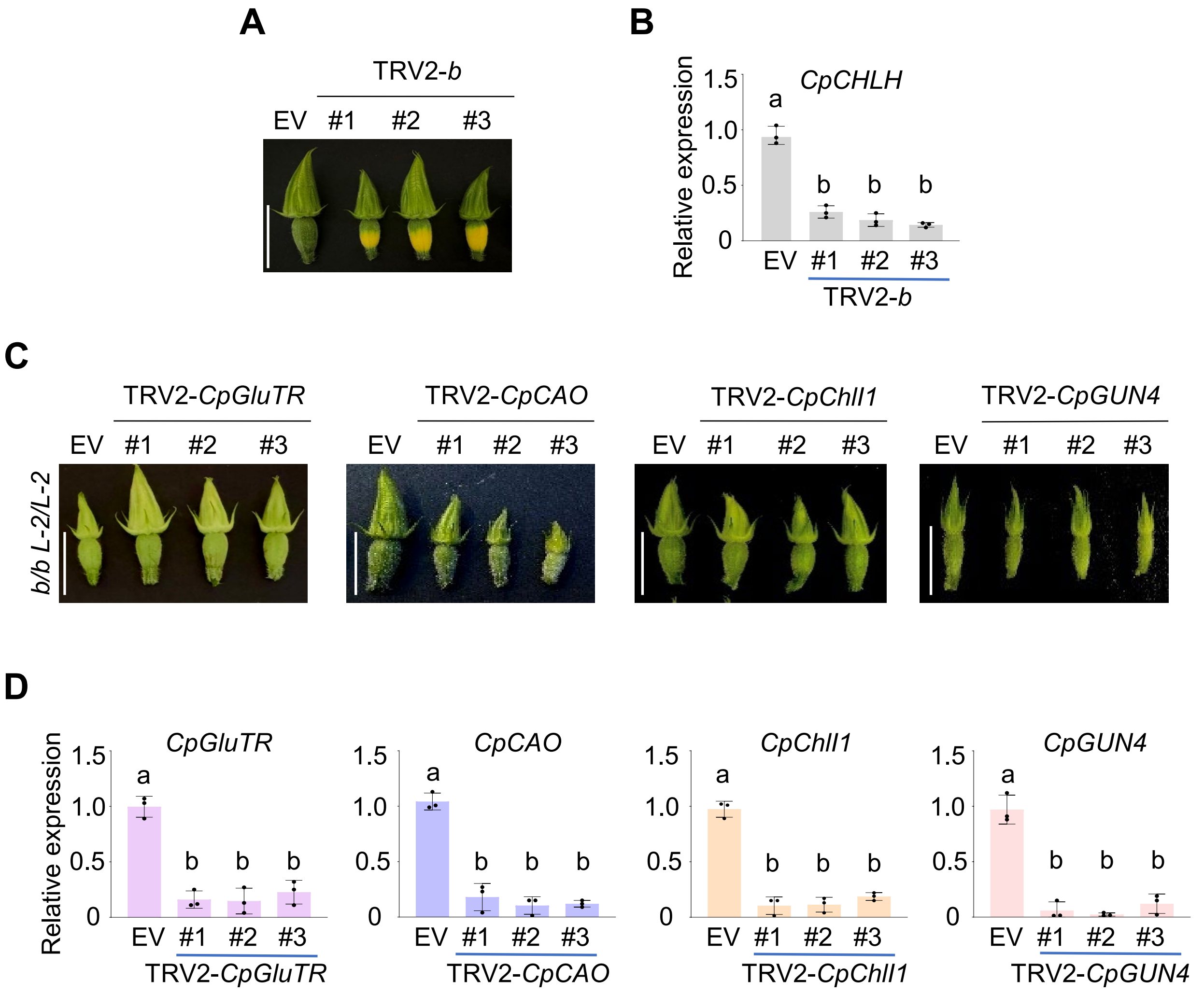


**Supplementary Figure S5.** Silencing of chlorophyll biosynthetic pathway genes by virus-induced gene silencing (VIGS). **A, C)** Fruit phenotype of TRV2-*b*, TRV2-*CpGluTR*, TRV2-*CpCAO*, TRV2-*CpChlI1* and TRV2-*CpGUN4* lines in *b/b L-2/L-2* genetic background at young fruit stage (< 5 day). Scale bar = 2 cm applicable to all images. **B, D)** RT-qPCR analysis of *CpChlH*, *CpGluTR*, *CpCAO*, *CpChlI1* and *CpGUN4* transcript levels in related VIGS lines vs EV (empty vector) controls in *b/b L-2/L-2* genetic background. Data in (**B, D**) are from three biological replicates and presented as mean ± SD (n = 3). Different letters denote significant differences as determined by one-way ANOVA followed by Tukey's multiple-comparison test ($p < 0.05$).

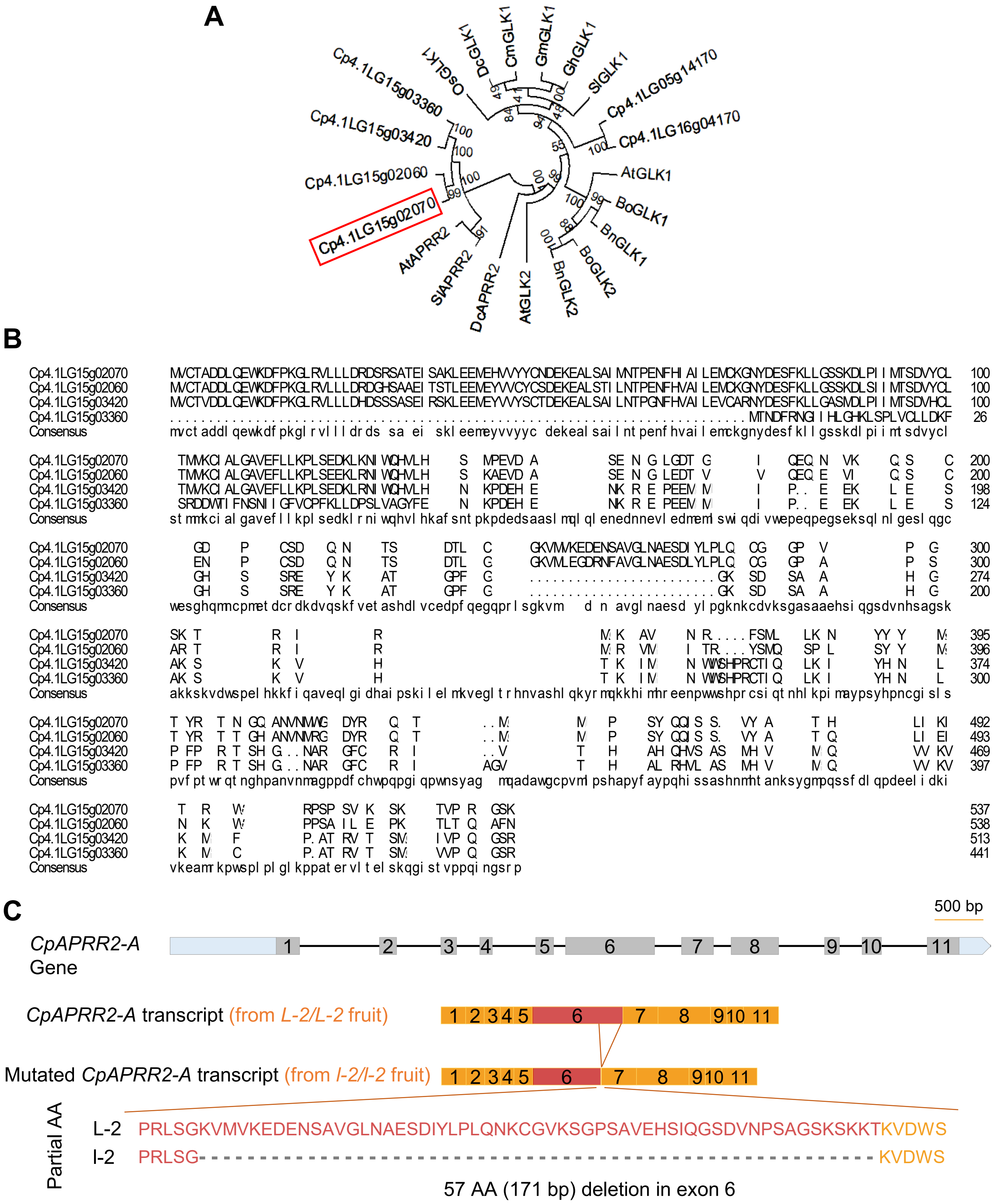


**Supplementary Figure S6.** APRR2 subfamily and *CpAPRR2-A* transcript structural variations from *L-2* and *l-2*. **A)** The amino acid sequence of Cp4.1LG05g02070 was retrieved from the *C. pepo* reference genome (Cucurbit Genomics Database), and its homologous protein sequences from other representative plants were collected through BLASTP searches on NCBI (https://blast.ncbi.nlm.nih.gov). Multiple sequence alignment was performed using M5b3 v5.0 software, and a phylogenetic tree was constructed using the neighbor-joining method with 1,000 bootstrap replicates. **B)** Full-length protein sequences of the four *C. pepo* APRR2 subfamily members were aligned using the DNAMAN v8.0 alignment software. **C)** *CpAPRR2-A* gene structure and its transcript from *L-2/L-2* and *l-2/l-2* genotype fruit. Numbered boxes indicate exons. The red boxes show the difference of *L-2* from *l-2* that produces a protein with 57 amino acid (AA) deletion.

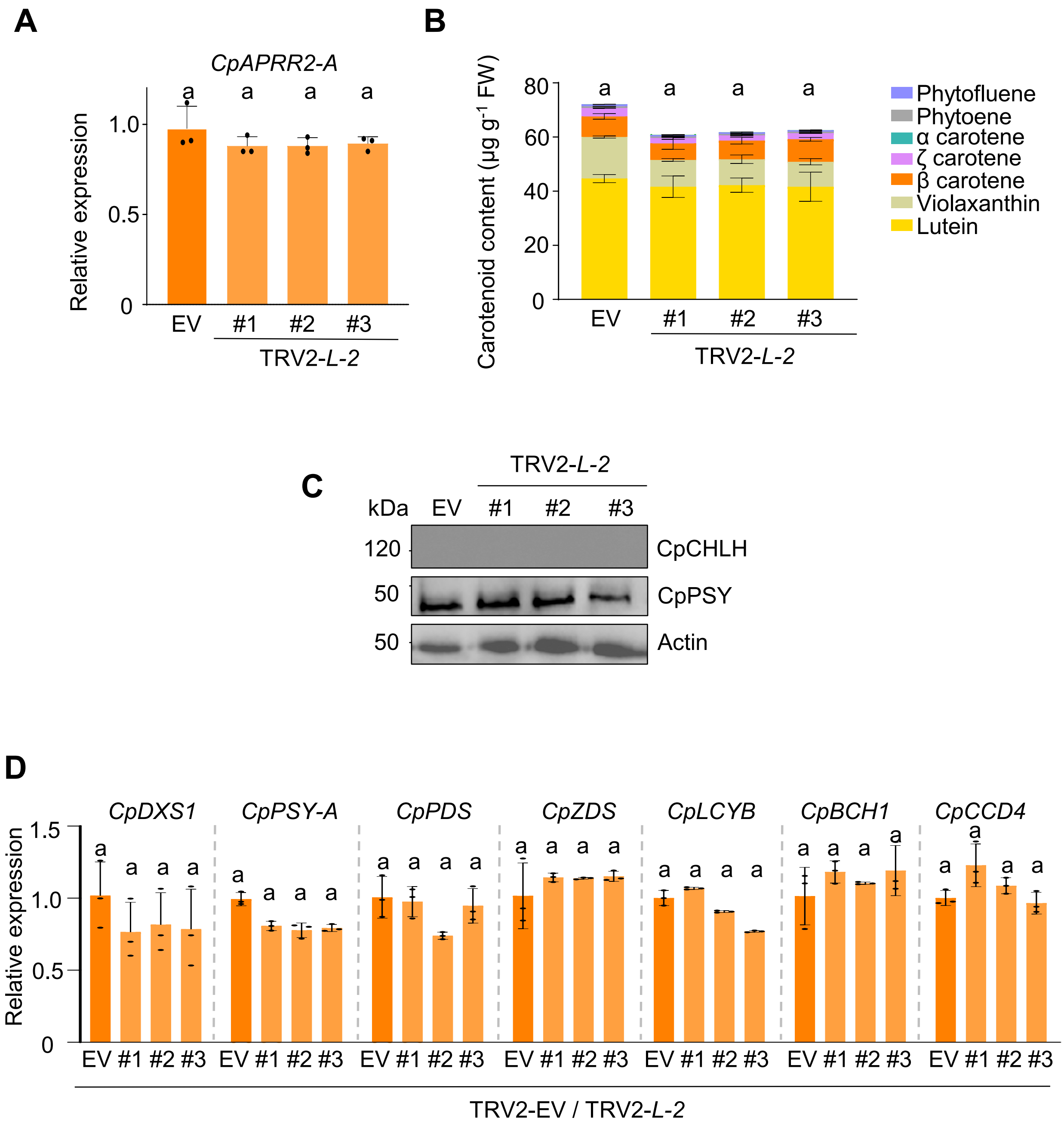


**Supplementary Figure S7.** Analysis of yellow sections of the TRV2-*L-2* fruit. **A)** The transcription level of *CpAPRR2-A* in yellow parts of empty vector (EV) control and TRV2-*L-2* fruit at 50 DAP stage in *B/B L-2/L-2* genotype. **B)** Carotenoid content and composition in yellow parts of EV and TRV2-*L-2* fruit. **C)** CpCHLH and CpPSY-A protein levels were detected by western blot in fruit (yellow parts) of TRV2-*L-2* lines at 50 DAP stage. Actin served as a loading control. **D)** Relative expression of carotenoid biosynthesis pathway genes were detected by RT-qPCR in yellow part of fruit at 50 DAP stage. Data in (**A**), (**B**), and (**D**) are from three biological replicates and presented as mean ± SD (n = 3). The same letters show no significant differences, as determined by one-way ANOVA followed by Tukey's multiple-comparison test ($p < 0.05$).

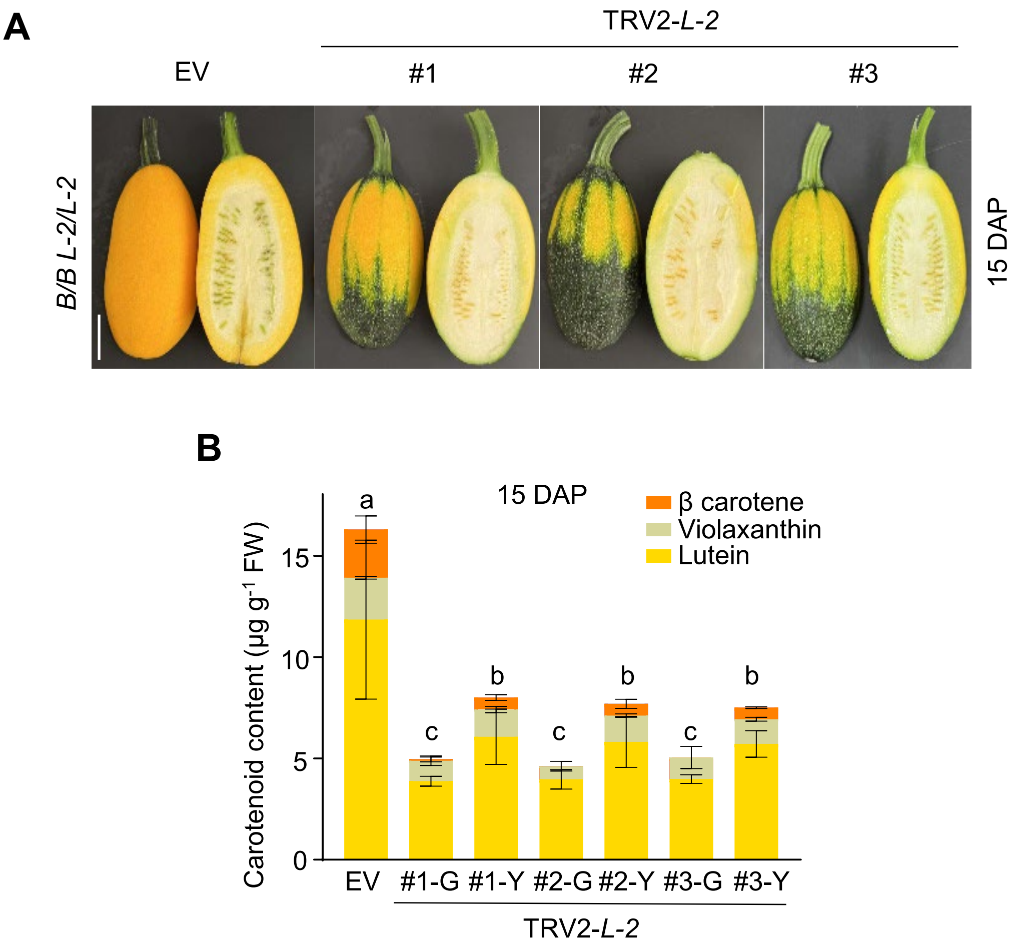


**Supplementary Figure S8.** TRV2-*L-2* silencing fruit at fruit developmental stage. **A)** Fruit phenotype of TRV2-*L-2* lines in *B/B L-2/L-2* background at fruit enlargement (15 DAP) stage. EV (empty vector) as control. Scale bar = 2 cm applicable to all images. **B)** Carotenoid content and composition at 15 DAP stage. G represents the green flesh part of the fruit, and Y represents the yellow part. Data in (**B**) are from three biological replicates and presented as mean ± SD (n = 3). Different letters denote significant differences, as determined by one-way ANOVA followed by Tukey's multiple-comparison test ($p < 0.05$).

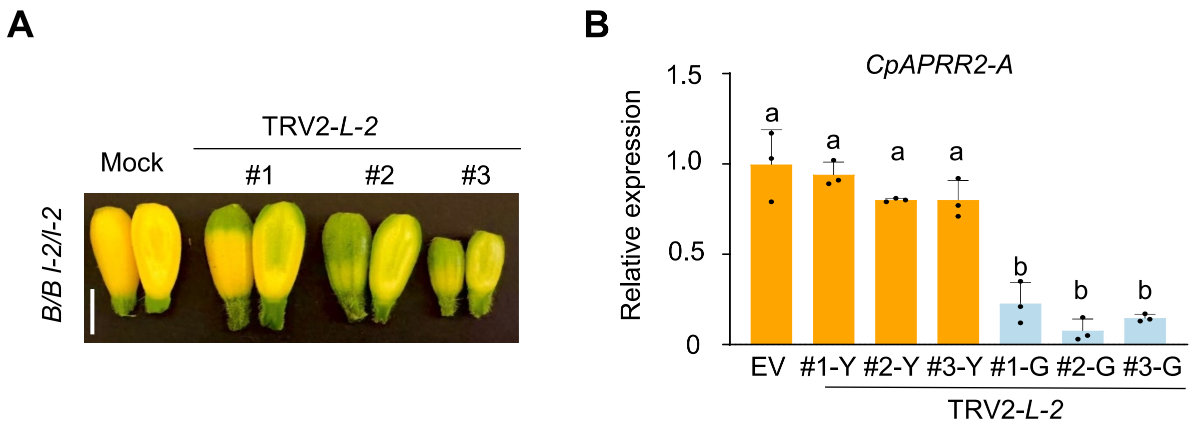


**Supplementary Figure S9.** Silencing of *l-2* in *B/B l-2/l-2* genotype. **A)** Fruit phenotype of TRV2-*l-2* in *B/B l-2/l-2* background at young (< 5 d) stage. EV (empty vector) as control. Scale bar = 1 cm applicable to all images. **B**) RT-qPCR analysis of *CpAPRR2-A* expression in yellow (Y) and green (G) parts of TRV2-*L-2* fruit at young stage. Data are from three biological replicates and presented as mean ± SD (n = 3). Different letters denote significant differences, as determined by one-way ANOVA followed by Tukey's multiple-comparison test ($p < 0.05$).

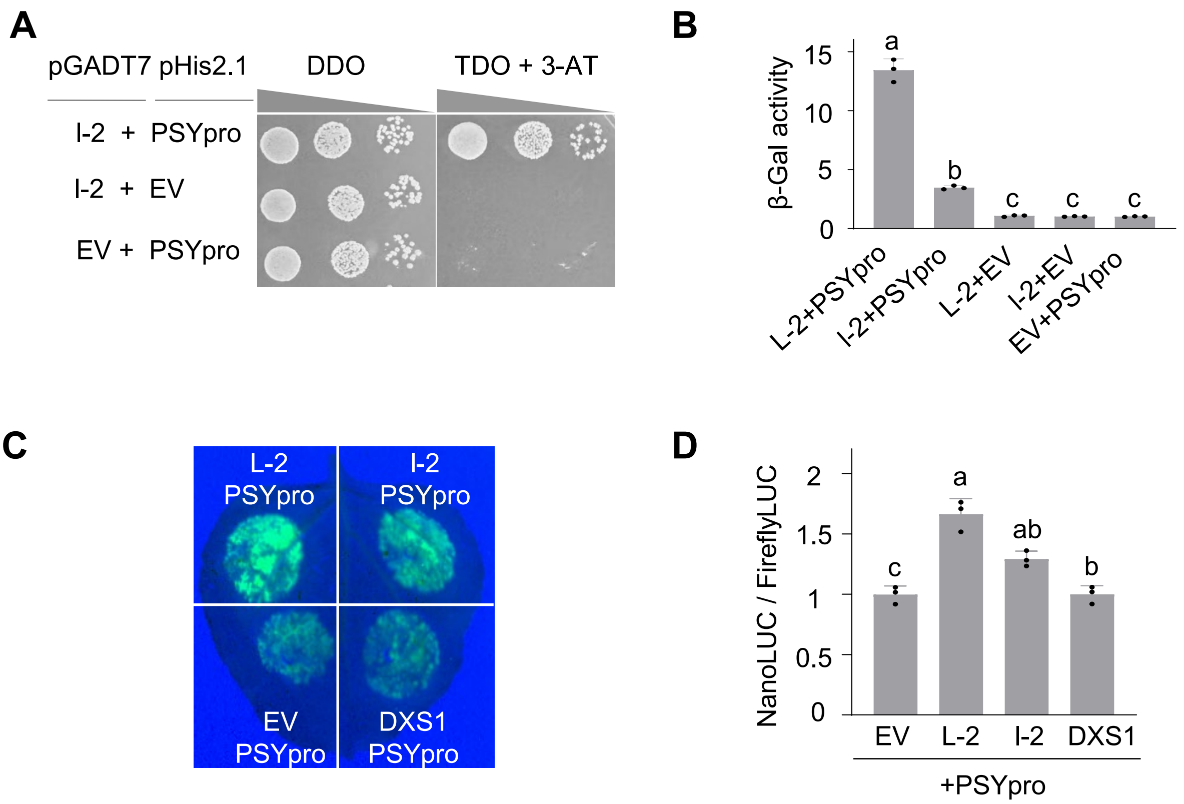


**Supplementary Figure S10.** *l-2* encoded mutated CpAPRR2-A with 57 amino acid deletion has reduced binding and transactivation activity. **A)** Y1H shows binding of *l-2* encoded protein (l-2) to *CpPSY-A* promoter (PSYpro). Yeast cells were co-transformed with the PSYpro-pHis2.1 construct and l-2-pGADT7 plasmid, then spread on DDO (double dropout medium, SD/–Leu/–Trp) and TDO (triple dropout medium, SD/–Leu/–Trp/–His with 80 mM 3-AT) selective medium. Empty-vector (EV) controls show no growth. **B)** Interaction strength of L-2 or l-2 with PSYpro and EV was assessed using a β-galactosidase reporter assay, and the activity was quantified in Miller units. **C)** Transactivation assay in *N. benthamiana* leaves showing differential activation of PSYpro by l-2. EV and DXS1 were used as negative control. **D)** Luciferase activity was quantified using protein extracts treated with luciferin or coelenterazine. Transactivation activity was normalized using FireflyLUC and expressed as NanoLUC/FireflyLUC. Data in (**B**) and (**D**) are from three biological replicates and presented as mean ± SD (n = 3). Different letters denote significant differences, as determined by one-way ANOVA followed by Tukey's multiple-comparison test ($p < 0.05$).

**Supplementary Table S1:** Carotenoid level and composition in flesh tissue of fruit at 50 DAP (μg $g^{-1}$ FW)

| Genotype | Phytoene | Phytofluene | ζ-Carotene | α-Carotene | β-Carotene | Violaxanthin | Lutein | Total carotenoids |
|---|---|---|---|---|---|---|---|---|
| *B/B L2/L2* | 0.77 ± 0.11 | 0.97 ± 0.14 | 3.72 ± 0.33 | 0.05 ± 0.09 | 7.93 ± 1.04 | 15.49 ± 2.55 | 49.37 ± 5.22 | 78.29 ± 3.21 |
| *B/B l2/l2* | 0 | 0 | 0 | 0 | 0.38 ± 0.15 | 0.84 ± 0.13 | 0.82 ± 0.19 | 2.04 ± 0.21 |
| *b/b L2/L2* | 0 | 0 | 0 | 0 | 0.23 ± 0.04 | 0.43 ± 0.12 | 1.25 ± 0.02 | 1.91 ± 0.05 |
| *b/b l2/l2* | 0 | 0 | 0 | 0 | 0.26 ± 0.04 | 0.82 ± 0.09 | 0.40 ± 0.03 | 1.48 ± 0.07 |

DAP: day after pollination; FW: fresh weight

**Supplementary Table S2:** Annotated genes in the candidate region of *B* locus

| Gene | Description |
|---|---|
| *Cp4.1LG10g11480* | Transmembrane protein |
| *Cp4.1LG10g11490* | Heat Stress Transcription Factor family protein |
| *Cp4.1LG10g11500* | Thioredoxin reductase |
| *Cp4.1LG10g11510* | ABC transporter A family member 1 |
| *Cp4.1LG10g11520* | ABC transporter A family member 1 isoform A |
| *Cp4.1LG10g11530* | ABC transporter A family member 1 |
| *Cp4.1LG10g11540* | Uncharacterized protein |
| *Cp4.1LG10g11550* | Leucine-rich repeat protein kinase family protein |
| *Cp4.1LG10g11560* | Magnesium chelatase H subunit |
| *Cp4.1LG10g11570* | L-lactate dehydrogenase 1 |
| *Cp4.1LG10g11580* | Exosome complex component RRP43 |
| *Cp4.1LG10g11590* | Exosome complex component RRP43 |
| *Cp4.1LG10g11600* | 40S ribosomal protein s2 |
| *Cp4.1LG10g11610* | tRNA-uridine aminocarboxypropyltransferase |
| *Cp4.1LG10g11620* | Protein of unknown function (DUF1138) |
| *Cp4.1LG10g11630* | Protein of unknown function, DUF642 |
| *Cp4.1LG10g11640* | Cellulose synthase |
| *Cp4.1LG10g11650* | Zinc finger AN1/C2H2 stress-associated protein 11-like |
| *Cp4.1LG10g11660* | Galactinol--sucrose galactosyltransferase |
| *Cp4.1LG10g11670* | Vvillin-2-like |
| *Cp4.1LG10g11680* | Unknown protein |
| *Cp4.1LG10g11730* | Calcium-dependent protein kinase |

**Supplementary Table S3:** Annotated genes in the candidate region of *L-2* locus

| Gene ID | Gene description |
|---|---|
| *Cp4.1LG05g02090* | Nitrate transporter 1.5 |
| *Cp4.1LG05g02020* | Pyruvate kinase |
| *Cp4.1LG05g02100* | Prolamin-like domain-containing protein |
| *Cp4.1LG05g01980* | Ribose-phosphate diphosphokinase |
| *Cp4.1LG05g01990* | Tobamovirus multiplication protein 2A-like |
| *Cp4.1LG05g02070* | Two-component response regulator-like APRR2 |
| *Cp4.1LG05g02060* | Two-component response regulator-like APRR2 |
| *Cp4.1LG05g02080* | Two-component response regulator-like APRR2 |
| *Cp4.1LG05g02030* | MSC domain-containing protein |

**Supplementary Table S4.** Primers used in this study

| Primer name | Primer sequence (5' - 3') | Usage |
|---|---|---|
| CpACTIN-qF | ttccgatgccctgaagttct | RT-qPCR |
| CpACTIN-qF | cattcggtcggcaataccag | RT-qPCR |
| AtACTIN-qF | ggttctacttaccgaggctcc | RT-qPCR |
| AtACTIN-qF | ggtaatcagtaaggtcacgac | RT-qPCR |
| DXS1-qF | acgtgtcgaaaactggtggt | RT-qPCR |
| DXS1-qR | cccgtttgtctgtagaatcctg | RT-qPCR |
| PSYA-qF | ggcaatatatgatgagcttgttga | RT-qPCR |
| PSYA-qR | gccttgcctcccatctatcc | RT-qPCR |
| PDS-qF | cgatatcctcgtcactggtcg | RT-qPCR |
| PDS-qR | ccaacatggactggttcgga | RT-qPCR |
| ZDS-qF | aaaggttggcgcagagaaga | RT-qPCR |
| ZDS-qR | cgccacagcatttcttgctt | RT-qPCR |
| LCYB-qF | tccgacttatcgtgacagcc | RT-qPCR |
| LCYB-qR | gcttcatatatcccaaatgaa | RT-qPCR |
| BCH1-qF | gtccctggtctctgcttcg | RT-qPCR |
| BCH1-qR | aacagcccgtacggaacacc | RT-qPCR |
| CCD4-qF | tgacggctcatccgaaatcc | RT-qPCR |
| CCD4-qR | atcggcacatcggactgttt | RT-qPCR |
| CHLH-qF | gttgggtttgcacccgattc | RT-qPCR |
| CHLH-qR | gcttcttctcagccctcttgt | RT-qPCR |
| CHLI1-qF | tcactctcttctccctcccc | RT-qPCR |
| CHLI1-qR | gaggcctttccttcttcccc | RT-qPCR |
| CHLD-qF | gctgctgatgctcctagacc | RT-qPCR |
| CHLD-qR | gaacaactgttgcgatgcca | RT-qPCR |
| GUN4-qF | aatcaacctcaacgcccact | RT-qPCR |
| GUN4-qR | ggaagtttttggctgcgagg | RT-qPCR |
| Cp4.1LG05g02090-qF | gcttggagatttgcaaccagaa | RT-qPCR |
| Cp4.1LG05g02090-qR | accccaaagaacgcaagagt | RT-qPCR |
| Cp4.1LG05g02020-qF | actttcagcgtcagatgggg | RT-qPCR |
| Cp4.1LG05g02020-qR | ctcggctctagttggtgtgg | RT-qPCR |
| Cp4.1LG05g02100-qF | gataaccgcggtggaaggat | RT-qPCR |
| Cp4.1LG05g02100-qR | atttgtccggcagtccatgt | RT-qPCR |
| Cp4.1LG05g01980-qF | tctccgtcccgaacgtgtta | RT-qPCR |
| Cp4.1LG05g01980-qR | catgcgagctgactccaaga | RT-qPCR |
| Cp4.1LG05g01990-qF | tcctggggatagaactggga | RT-qPCR |
| Cp4.1LG05g01990-qR | accacaagggccaacaagaa | RT-qPCR |
| Cp4.1LG05g02070-qF | ggatgcaagctgatgcatgg | RT-qPCR |
| Cp4.1LG05g02070-qR | gggcgtgccatagctcttat | RT-qPCR |
| Cp4.1LG05g02060-qF | tccaaggcaaaaactagccga | RT-qPCR |
| Cp4.1LG05g02060-qR | gccgtcccgatcaaggagaa | RT-qPCR |
| Cp4.1LG05g03360-qF | ttgacggtgatgggagtcag | RT-qPCR |
| Cp4.1LG05g03360-qR | agaacgaaggtgccggaga | RT-qPCR |

| | | |
|---|---|---|
| Cp4.1LG05g03420-qF | gctcaccaatcccaagatgttt | RT-qPCR |
| Cp4.1LG05g03420-qR | catccattccctcccctgtg | RT-qPCR |
| Cp4.1LG05g04170-qF | ggtcaggtcagcaacatgga | RT-qPCR |
| Cp4.1LG05g04170-qR | ataagcccctcaaagcctgc | RT-qPCR |
| AtGUN5-qF | tcttcacacagacgaacccg | RT-qPCR |
| AtGUN5-qR | tgttgagagattgcacggct | RT-qPCR |
| AtPSY-qF | tctattgtggctcttgtttggtg | RT-qPCR |
| AtPSY-qR | cgaaaggacgaccacggaaa | RT-qPCR |
| At4G03770-chip-actin-qF | ggctcgtgcggaaatcattc | Chip-qPCR |
| At4G03770-chip-actin-qR | aattgttggcgcatgccttt | Chip-qPCR |
| PSYpro-ARE1-qF | attcaataaagtcgctaatt | Chip-qPCR |
| PSYpro-ARE1-qR | caaactaaggcaatttttac | Chip-qPCR |
| PSYpro-ARE2-qF | ggttcgtttctcctccctcc | Chip-qPCR |
| PSYpro-ARE2-qR | ctccgtgaccaacaaacacc | Chip-qPCR |
| PSYpro-ARE3-qF | ataatcgggttcttatatta | Chip-qPCR |
| PSYpro-ARE3-qR | aattggaaaatgaagtggaa | Chip-qPCR |
| PSYpro-ARE4-qF | aacttgctttttccacttca | Chip-qPCR |
| PSYpro-ARE4-qR | atcaatcgattttcaaaatt | Chip-qPCR |
| PSYpro-negative control-qF | aggtgtttgttggtcacgga | Chip-qPCR |
| PSYpro-negative control-qR | tagcgatgctttgacacgga | Chip-qPCR |
| GluTR-qF | tacaagccgctcaccgaaat | RT-qPCR |
| GluTR-qR | gatgcatgctccgacgtttc | RT-qPCR |
| CAO-qF | cacctcggttctgtcaacga | RT-qPCR |
| CAO-qR | ccaggccaaacccagatcat | RT-qPCR |
| TRV2-b-pDONR207-F | ggggacaagtttgtacaaaaaagcaggcttaatggcgtccttgatgtcatcg | VIGS |
| TRV2-b-pDONR207-R | ggggaccactttgtacaagaaagctgggtcgcgtcgagtcgatccctttct | VIGS |
| TRV2-L-2-pDONR207-F | ggggacaagtttgtacaaaaaagcaggcttagaggacaaactgaagaatatc | VIGS |
| TRV2-L-2-pDONR207-R | ggggaccactttgtacaagaaagctgggtccacatctttgtcactgcaatc | VIGS |
| TRV2-GluTR-pDONR207-F | ggggacaagtttgtacaaaaaagcaggcttaatgcatcttcaagcacaaact | VIGS |
| TRV2-GluTR-pDONR207-R | ggggaccactttgtacaagaaagctgggtcgtcaacagcgaatcgaacgcc | VIGS |
| TRV2-CAO-pDONR207-F | ggggacaagtttgtacaaaaaagcaggcttagaaacgctatttgacgtggaa | VIGS |
| TRV2-CAO-pDONR207-R | ggggaccactttgtacaagaaagctgggtccgcagtggcaacccgtgcttc | VIGS |
| TRV2-CHLI1-pDONR207-F | ggggacaagtttgtacaaaaaagcaggcttacaaagagttccgtgaatctta | VIGS |
| TRV2-CHLI1-pDONR207-R | ggggaccactttgtacaagaaagctgggtcccagaatcaatagattccaat | VIGS |
| TRV2-GUN4-pDONR207-F | ggggacaagtttgtacaaaaaagcaggcttacctcaaatcaacctcaacgcc | VIGS |
| TRV2-GUN4-pDONR207-R | ggggaccactttgtacaagaaagctgggtcccatcgctgtatttctgccag | VIGS |
| pDEST32-L-2-F | ggggacaagtttgtacaaaaaagcaggcttaatggtttgcactgccgacgatttacaa | Transactiviation assay |
| pDEST32-L-2-R | ggggaccactttgtacaagaaagctgggtcgggaggtttggagccgttgattcgagg | Transactiviation assay |
| pDEST32-l-2-F | ggggacaagtttgtacaaaaaagcaggcttaatggtttgcactgccgacgatttacaa | Transactiviation assay |
| pDEST32-l-2-R | ggggaccactttgtacaagaaagctgggtcgggaggtttggagccgttgattcgagg | Transactiviation assay |
| pDUAL-UBQ-PSYpro-p207-F | ggggacaagtttgtacaaaaaagcaggcttagatatcaaaagatttaaaaaatagt | Transactiviation assay |
| pDUAL-UBQ-PSYpro-p207-R | ggggaccactttgtacaagaaagctgggtcatcaatcgattttcaaaatttcact | Transactiviation assay |
| pGWB17-L-2-pDONR207-F | ggggacaagtttgtacaaaaaagcaggcttaatggtttgcactgccgacgatttacaa | Overexpression lines |
| pGWB17-L-2-pDONR207-R | ggggaccactttgtacaagaaagctgggtcgggaggtttggagccgttgattcgagg | Overexpression lines |

| | | |
|---|---|---|
| pGWB17-l-2-pDONR207-F | ggggacaagtttgtacaaaaaagcaggcttaatggtttgcactgccgacgatttacaa | Overexpression lines |
| pGWB17-l-2-pDONR207-R | ggggaccactttgtacaagaaagctgggtcgggaggtttggagccgttgattcgagg | Overexpression lines |
| pGWB17-B-pDONR207-F | ggggacaagtttgtacaaaaaagcaggcttaatggcgtccttgatgtcatcg | Overexpression lines |
| pGWB17-B-pDONR207-R | ggggaccactttgtacaagaaagctgggtctcgatcgaccccctcgatctt | Overexpression lines |
| pGWB17-b-pDONR207-F | ggggacaagtttgtacaaaaaagcaggcttaatggcgtccttgatgtcatcg | Overexpression lines |
| pGWB17-b-pDONR207-R | ggggaccactttgtacaagaaagctgggtctcgatcgaccccctcgatctt | Overexpression lines |
| pGADT7-L-2-F | cgcgggatccatggtttgcactgccgacgatttacaa | Y1H |
| pGADT7-L-2-R | ggactcgaggggaggtttggagccgttgattcgagg | Y1H |
| pGADT7-l-2-F | cgcgggatccatggtttgcactgccgacgatttacaa | Y1H |
| pGADT7-l-2-R | ggactcgaggggaggtttggagccgttgattcgagg | Y1H |
| pHis2.1-PSYpro-F | ccggaattcgatatcaaaagatttaaaaaatagta | Y1H |
| pHis2.1-PSYpro-R | cggactagtatcaatcgattttcaaaatttcactg | Y1H |
| Nub-b-pDONR207-F | ggggacaagtttgtacaaaaaagcaggcttaatggcgtccttgatgtcatcg | Y2H |
| Nub-b-pDONR207-R | ggggaccactttgtacaagaaagctgggtctcgatcgaccccctcgatctt | Y2H |
| Nub-B-pDONR207-F | ggggacaagtttgtacaaaaaagcaggcttaatggcgtccttgatgtcatcg | Y2H |
| Nub-B-pDONR207-R | ggggaccactttgtacaagaaagctgggtctcgatcgaccccctcgatctt | Y2H |
| Cub-L-2-pDONR207-F | ggggacaagtttgtacaaaaaagcaggcttaatggtttgcactgccgacgatttacaa | Y2H |
| Cub-L-2-pDONR207-R | ggggaccactttgtacaagaaagctgggtcgggaggtttggagccgttgattcgagg | Y2H |
| nYFP-L-2-pDONR207-F | ggggacaagtttgtacaaaaaagcaggcttaatggtttgcactgccgacgatttacaa | BiFC |
| nYFP-L-2-pDONR207-R | ggggaccactttgtacaagaaagctgggtcgggaggtttggagccgttgattcgagg | BiFC |
| cYFP-b-pDONR207-F | ggggacaagtttgtacaaaaaagcaggcttaatggcgtccttgatgtcatcg | BiFC |
| cYFP-b-pDONR207-R | ggggaccactttgtacaagaaagctgggtctcgatcgaccccctcgatctt | BiFC |
| cYFP-B-pDONR207-F | ggggacaagtttgtacaaaaaagcaggcttaatggcgtccttgatgtcatcg | BiFC |
| cYFP-B-pDONR207-R | ggggaccactttgtacaagaaagctgggtctcgatcgaccccctcgatctt | BiFC |
| LM1-102611-F | tgctaatgtgcaccacccaa | Low resolution marker |
| LM1-102611-R | cgacgaaatgtgaagtgggc | Low resolution marker |
| LM2-697458-F | caaatcgtcgcccgcaaatc | Low resolution marker |
| LM2-697458-R | ggctttgctttgggcttctc | Low resolution marker |
| LM3-870183-F | acagacagagcacaggcaaa | Low resolution marker |
| LM3-870183-R | tatgcagctttctgactcgca | Low resolution marker |
| LM4-1109841-F | acatccaccactcggattgt | Low resolution marker |
| LM4-1109841-R | cggtcttcattacaggtttgcg | Low resolution marker |
| LM5-1606591-F | aggttgtagggtataggtcggt | Low resolution marker |
| LM5-1606591-R | tgaaccctcatgtcgtagca | Low resolution marker |
| LM6-1645151-F | cctgtttgactccatcctcaaat | Low resolution marker |
| LM6-1645151-R | actggtttgatctgggaggt | Low resolution marker |
| LM7-1752280-F | tgcaacaagcaagaagtggg | Low resolution marker |
| LM7-1752280-R | cgcatcaaaacgcagtcagg | Low resolution marker |
| HM1-697458-F | caaatcgtcgcccgcaaatc | High resolution marker |
| HM1-697458-R | ggctttgctttgggcttctc | High resolution marker |
| HM2-961548-F | ggttccactttgaaggccc | High resolution marker |
| HM2-961548-R | ggaggaggaccaagcaatcc | High resolution marker |
| HM3-1054822-F | gaaagtcgtgagacaagtggt | High resolution marker |
| HM3-1054822-R | cgttaagcccatgttttggca | High resolution marker |

| | | |
|---|---|---|
| HM4-1109841-F | acatccaccactcggattgt | High resolution marker |
| HM4-1109841-R | cggtcttcattacaggtttgcg | High resolution marker |
| HM5-1247457-F | tggtagtggttgagtggtgc | High resolution marker |
| HM5-1247457-R | acataacgggtcaaagcagaca | High resolution marker |
| HM6-1388782-F | agccctacggtttagggtca | High resolution marker |
| HM6-1388782-R | tcggcccaattgtaaaccca | High resolution marker |
| HM7-1606591-F | aggttgtagggtataggtcggt | High resolution marker |
| HM7-1606591-R | tgaaccctcatgtcgtagca | High resolution marker |
